# Decoding the peripheral transcriptomic and meta-genomic response to music in Autism Spectrum Disorder *via* saliva-based RNA sequencing

**DOI:** 10.1101/2025.07.04.663204

**Authors:** Andrea Cavenaghi, Nour El Zahraa Mallah, Laura Navarro, Group Sensogenomics Working, Federico Martinón-Torres, Alberto Gómez-Carballa, Antonio Salas

**Affiliations:** Unidade de Xenética, Instituto de Ciencias Forenses, Facultade de Medicina, Universidade de Santiago de Compostela, and Genética de Poblaciones en Biomedicina (GenPoB) Research Group, Instituto de Investigación Sanitaria (IDIS), 15706 Hospital Clínico Universitario de Santiago (SERGAS), Galicia, Spain; Genetics, Vaccines and Infections Research Group (GenViP), Instituto de Investigación Sanitaria de Santiago, 15706 Universidade de Santiago de Compostela, Santiago de Compostela, Galicia, Spain; Centro de Investigación Biomédica en Red de Enfermedades Respiratorias (CIBER-ES), Madrid, Spain; Translational Pediatrics and Infectious Diseases, Department of Pediatrics, 15706 Hospital Clínico Universitario de Santiago de Compostela, Santiago de Compostela, Galicia, Spain

**Keywords:** ASD, Transcriptomics, Gene Expression, RNA-seq

## Abstract

Saliva-based RNA sequencing (RNA-seq) poses technical challenges due to high bacterial content, RNA degradation, and sample heterogeneity. This study investigates the transcriptional effects of music exposure in individuals with autism spectrum disorder (ASD) using this non-invasive approach. To address saliva-specific limitations, we employed two complementary library preparation methods, Poly-A selection and Human-Enriched protocols, allowing us to maximize human transcript detection and ensure reproducibility. By merging them, we ensured reproducibility and captured both host and microbial signals. While each dataset individually revealed a limited number of differentially expressed genes (DEGs), their integration enhanced biological resolution. Among the consistently modulated genes were *HERC6*, *TSPAN5*, and *REM2*, pointing to music-induced transcriptional changes relevant to neurodevelopmental and immune processes. Functional enrichment analyses highlighted pathways involved in immune regulation, oxidative phosphorylation, and epithelial differentiation. These findings align with evidence of immune dysregulation, mitochondrial dysfunction, and altered cellular communication in ASD. Importantly, co-expression network analysis identified modules significantly correlated with music exposure. Notably, the *AKNA* module, previously associated with ASD risk, was downregulated and enriched for Ras-related GTPase signaling and immune pathways, suggesting that music may modulate intracellular signaling and inflammation. Conversely, upregulation of the *UBE2D3* module pointed to activation of endoplasmic reticulum stress response mechanisms, a known contributor to ASD neurodevelopmental deficits. These results suggest that music engages specific stress-adaptive and immunomodulatory networks in buccal cells, potentially reflecting systemic effects. Our exploratory metagenomic analysis highlights 15 microbial species with consistent abundance shifts across both methods. Notably, *Acidipropionibacterium acidipropionici* and *Propionibacterium freudenreichii*, associated with propionic acid production, emerged as music-responsive taxa. Elevated propionic acid has been implicated in ASD-like behaviors and neuroinflammation, suggesting a microbiota-mediated pathway. Music may influence both host gene expression and oral microbiota, potentially affecting neuroimmune processes via the microbiota–brain axis. Although exploratory, the results support the feasibility of using saliva for integrated molecular profiling in ASD.

## Introduction

Autism Spectrum Disorder (ASD) is a neurodevelopmental condition involving deficits in communication, behavior, and motor skills ([1–3]; https://icd.who.int/). Symptoms typically emerge between 18 and 24 months, and diagnosis is based on persistent social interaction difficulties, communication challenges, and repetitive behaviors [4]. ASD affects about 1.68% of U.S. children and 1.4% in Europe, with a male-to-female ratio of 3.5:1 [5]. This disparity may be partially due to underdiagnosis of females, who often exhibit subtler symptoms and camouflaging behaviors, suggesting that females may need a higher genetic or environmental threshold to express ASD traits [6]. However, when diagnosed, females often present with more severe symptoms and intellectual disability [7]. ASD spans a wide spectrum, from severe impairments to mild social differences [8], and some individuals also show developmental regression, particularly in language and social skills [9]. Educational, occupational, and social difficulties are frequent. However, others can exhibit exceptional abilities, such as enhanced pitch recognition or visual processing [10–13]. Comorbid conditions are common, including intellectual disability (∼31%), sleep disturbances, seizures, anxiety, Attention-Deficit/Hyperactivity Disorder (ADHD), Obsessive-Compulsive Disorder (OCD), and mood disorders [14, 15]. Needs vary; some people live independently, while others require lifelong support [16].

ASD’s causes are multifactorial, involving genetic and environmental components. Brain abnormalities such as cerebellar malformations and early overgrowth are being studied as potential causal factors [17, 18]. Genetic research implicates genes involved in brain development and synaptic function [19–21]. Environmental risk factors include prenatal exposure to valproic acid or thalidomide, older parental age, low birth weight, and cesarean delivery [22]. Maternal inflammation during pregnancy is another risk factor for ASD. Cytokines like IL-6 and IL-17a, elevated during infections, may disrupt fetal brain development and promote ASD traits [23]. Postnatal immune profiles also differ in ASD individuals, implicating immune dysregulation. By contrast, folic acid supplementation during pregnancy appears to have a protective effect [22].

On the other hand, music engages widespread brain regions and influences emotion, cognition, and motor skills [24]. It stimulates dopamine release, affects autonomic responses, and promotes neuroplasticity. Its effects vary by age, sex, culture, and musical training [24, 25]. Music therapy is used in diverse clinical contexts such as addiction, Post-Traumatic Stress Disorder (PTSD), dementia, and brain injury [26]. In ASD, music is promising due to the usually higher auditory sensitivity and pitch memory of the patients [11]. While auditory sensitivity can lead to overload in ASD patients, it also allows deep engagement with music’s predictability and structure [27]. Active and passive music therapy can support not only emotional regulation and social communication but also improve emotional processing, motivation, attention, and interaction [28]. Structured programs, in turn, strengthen parent-child relationships while reducing symptom severity [29]. Music stimulation increases oxytocin and vasopressin, neuropeptides involved in empathy and bonding. Oxytocin enhances social cue recognition and has been genetically linked to ASD [30]. Compared to speech therapy or medication, music therapy often yields greater improvements in life quality [31]. Neuroimaging studies have shown that music therapy enhances brain connectivity, especially between auditory and motor regions [32]. However, most research studies focused on children and lacked standardization. Given individual variability in music sensitivity, personalized approaches are essential. In this context, the emerging field of sensogenomics investigates how sensory stimuli like music influence gene expression and brain function, offering new insights into ASD.

Many children with ASD exhibit significant fear and avoidance of medical procedures, particularly those involving needles [33, 34] and, therefore, alternative less invasive approaches are under investigation. Though exposure therapy may help reduce stress related to blood draws, its application is limited by safety and logistical constraints [35]. In this context, saliva presents a non-invasive, easily collectable alternative for biomarker discovery, supplying DNA, RNA, and proteins for molecular analysis, including identification of polymorphisms linked to ASD. Saliva is increasingly recognized as a valuable non-invasive biofluid for biomedical research and biomarker discovery. In the context of neurological conditions, saliva is being explored for monitoring brain health [36, 37]. Parasympathetic and sympathetic stimulation of salivary glands triggers acetylcholine and noradrenaline release to the buccal cavity, respectively, promoting protein secretion (**Figure 1A**); these neurotransmitters have been detected in rodent salivary glands [38–40]. The nervous regulation of the salivary glands supports the concept of a bidirectional oral-brain axis, where local inflammatory stages may influence each other [41].

**Figure 1.**
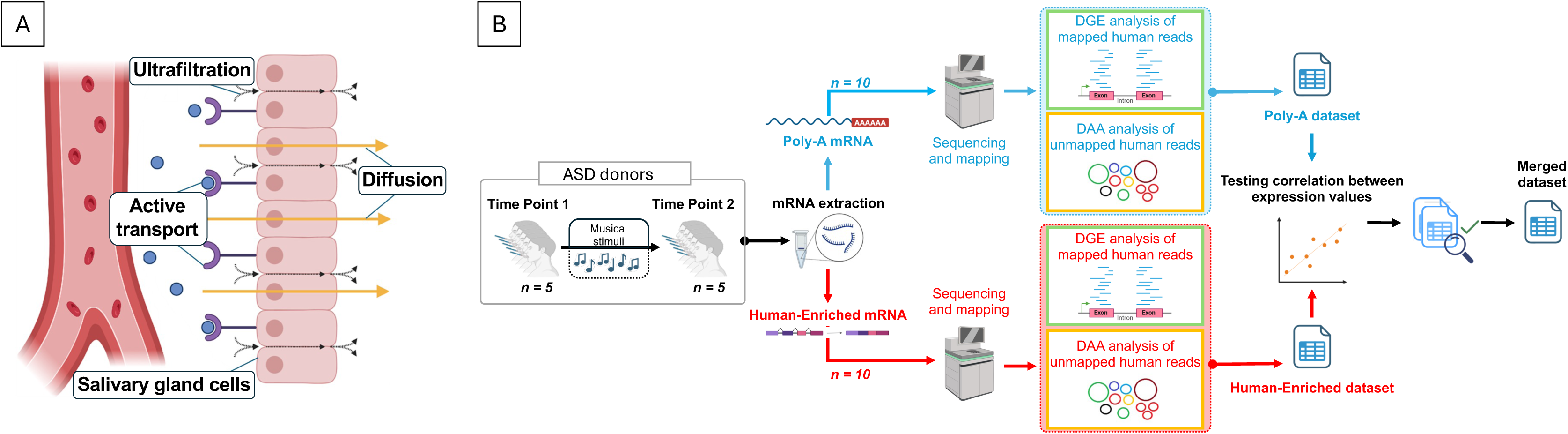
Complementary schemes illustrating the conceptual framework of this study. (A) Illustration of the transportation of biomolecules from blood capillaries to saliva (adapted from [57]). (B) Graphic scheme showing the design of the present work. The figure was built using Biorender resources (https://biorender.com/).

In the context of ASD, small RNA molecules such as microRNAs (miRNAs), piRNAs, and snoRNAs are involved in neurodevelopment and synaptic regulation. In a 2018 study, 32 RNA markers in saliva distinguished ASD from controls with 85% accuracy. Changes in miRNAs like *miR-146b-5p* and *piR-6463* were found diagnostically relevant [42]. *miR-141-3p* expression was found to be negatively correlated with the oral bacteria *Tannerella* in ASD children’s saliva, suggesting that dysregulation of miRNAs and the oral microbiome dysbiosis may be linked to cognitive impairments [43]. Indeed, 70% of ASD children have gastrointestinal symptoms [44]. Microbiome-related inflammation could impair the blood-brain barrier, affecting brain function. Hormonal markers in saliva from ASD patients, such as cortisol, have shown different results [45, 46]. Studies in different tissues, including saliva, have shown that oxytocin levels are lower in ASD children [47]. Several studies have tried to “treat” ASD social impairments with oxytocin treatments: some of them showed improvements [48, 49], while others showed no differences between the oxytocin treatment and the placebo treatment [50, 51].

Despite its advantages, saliva-based diagnostics face technical challenges. Only about 20% of RNA sequences align with the human genome, while 30% match known oral microbes, leaving 50% unclassified, likely representing fungi, viruses, or degraded fragments [52, 53]. Human RNA is easily degraded, and bacteria produce abundant RNA. In datasets like RNAgE-WT, human RNA comprised just 0.1% of reads, while bacterial RNA exceeded 75% [54]. Low analyte concentrations, sample variability, and degradation during handling (e.g., freeze-thaw cycles) further complicate analysis [55–57].

Exploring music’s effect on gene expression in ASD through saliva offers a promising, non-invasive research avenue. However, it involves two major obstacles: the complexity of human *vs.* microbial RNA content, and the novelty of assessing transcriptional changes induced by music in a population with neurobiological heterogeneity. Addressing these challenges could advance personalized music-based therapies.

This pilot study evaluated the feasibility of using saliva to analyze human RNA changes before and after music exposure in individuals with ASD. We employed two complementary methods of RNA-seq library preparation, and the results revealed reproducible transcriptomic responses, supporting saliva’s potential as a non-invasive tool for exploring the neurobiological impact of music in ASD. These findings lay the groundwork for future research into personalized, music-based interventions for neurodevelopmental disorders.

## Material and methods

### Participants and sample collection

This study was conducted as part of the Sensogenomics project (https://sensogenomics.com; [58–60]), which explores the biomolecular and physiological effects of music in diverse population groups. An experimental cinematic music concert was held on September 27^th^, 2024, at the Auditorio de Galicia in Santiago de Compostela (Galicia, Spain) with the Real Filharmonía de Galicia (https://www.rfgalicia.org). The event was organized in collaboration with the “Banda Municipal de Música de Santiago de Compostela” and led by conductor Baldur Brönnimann under his artistic direction. The concert had a total duration of 60 minutes and was divided into two parts. To minimize response bias, the musical program was not disclosed in advance. All participants remained seated throughout the performance to eliminate the influence of physical activity. The environment was carefully designed to be comfortable, calm, and non-invasive, thereby reducing stress or other external factors that could affect physiological responses. The audience consisted of individuals from distinct groups, including individuals diagnosed with ASD, healthy control participants, and general spectators. For the present pilot study, a subset of five individuals with ASD was selected. The mean age of the ASD subjects was 23.8 years (8-37 years), and 60% were females. To ensure the reliability and comparability of transcriptomic data, all participants were instructed to abstain from eating, drinking, smoking, chewing gum, or engaging in physical activity for at least 30 minutes before saliva collection.

Saliva samples were collected immediately before and after the concert, simultaneously for all participants. The collection was performed under the supervision of trained healthcare professionals to guarantee consistency and adherence to the protocol. Each participant used an Oragene RNA saliva collection device (ORE-100, DNA Genotek), which is optimized for the non-invasive collection, stabilization, and transport of high-quality RNA from saliva. Participants were instructed to rub the sponge tips of the device along both sides of their gums, avoiding contact with teeth, until sufficient saliva was absorbed. The sponge tips were then inserted into tubes containing 1 mL of stabilizing solution, which preserves RNA integrity. Samples were subsequently stored at room temperature until processing.

Written informed consent was obtained from the legal guardians or responsible parties of all participating individuals. The study was approved by the Ethics Committee of the Xunta de Galicia (registration code: 2020/021) and conducted in accordance with the principles outlined in the Declaration of Helsinki. **Figure 1B** shows a scheme of the experimental design used in the present study.

### RNA isolation and RNA-sequencing

RNA was isolated from 500□μL of saliva using the RNeasy Micro Kit (Qiagen), using the protocol provided by the extraction kit as recommended by the Oragene tubes supplier. Samples were subjected to a DNase treatment to remove DNA. Further purification, concentration, and DNAse treatment were performed using the RNA Clean & Concentrator Kit (Zymo). Final RNA concentration was verified with Nanodrop, yielding >50□ng/μL, suitable for downstream applications.

RNA quantity and integrity (RIN) were assessed using the TapeStation 4200 system (Agilent), and RNA concentration was quantified using the Qubit fluorometer (Thermo Fisher Scientific). Two different Illumina library preparation strategies were employed: *i*) Illumina RNA Prep with Enrichment (https://emea.illumina.com/products/by-type/sequencing-kits/library-prep-kits/rna-prep-enrichment.html), which uses hybrid capture to enrich for specific RNA targets, including non-coding RNAs, degraded RNA, or low-input samples (hereafter referred to as the Human-Enriched library), and *ii*) TruSeq Stranded mRNA, which selectively captures polyadenylated mRNA (poly-A selection), primarily targeting coding transcripts and thereby focusing on protein-coding gene expression (hereafter referred to as the Poly-A library). Human-Enriched libraries were prepared using the Illumina RNA Prep with Enrichment (ref. 20040537) workflow, which includes RNA fragmentation, reverse transcription into cDNA, adapter ligation, library amplification, purification, and target enrichment via hybridization capture using the Illumina Exome Panel (ref. 20020183). For the Poly-A library, strand-specific libraries were generated using the TruSeq Stranded mRNA technology (ref. 20020594). Indexing was performed using unique dual indexes with the IDT for Illumina TruSeq RNA UD Indexes kit (ref. 20022371).

Sequencing for both library types was performed on the NovaSeq 6000 System (Illumina) using a paired-end configuration (2×100 bp). Each sample generated approximately 30 million paired-end reads when using the Human-Enriched and 60 million paired-end reads with the Poly-A, with an average of 85% of bases achieving a quality score above Q30.

### Transcriptomic data processing and statistical analysis

To assess the quality of the raw reads, *FastQC* (v0.11.9) and *MultiQC* were employed. Following the initial quality assessment, reads were processed using *Trimmomatic* [61] to remove low-quality bases and residual adapter sequences. Reads were further filtered using the sliding window approach, where sequences are scanned from the 5’ to 3’ end and trimmed when the average quality within a defined window falls below a threshold. High-quality reads were aligned to the *Homo sapiens* reference genome (GRCh38/hg38) using *STAR* (Spliced Transcripts Alignment to a Reference) version 2.7.9a [62]. Gene-level quantification was performed using the *HTSeq* package (version 2.0.3) [63]. Data normalization and differential expression (DE) analysis was conducted using the *DESeq2* package (version 1.46.0) in RStudio (v4.4.2) [64]. To evaluate the robustness and reproducibility of the results, two datasets derived from different RNA isolation methods, Human-Enriched and Poly-A selected libraries, were compared. Exploratory data visualization was performed using histograms of read counts, Principal Component Analysis (PCA), hierarchical clustering heatmaps, and volcano plots. These analyses were executed with R packages *ggplot2* [65], *ComplexHeatmap* [66], and *EnhancedVolcano* [67]. *DESeq2* package was used to detect differentially expressed genes (DEGs) after the musical stimuli when compared to the baseline. We include the patient ID in the model as a random effect to account for patient-to-patient differences in the baseline. Genes common to both datasets and displaying statistically significant differential expression (*P*-value < 0.05) were examined *via* Pearson correlation to assess consistency across the two preparation methods. Upon confirming a strong correlation, a merged dataset was constructed by summing the read counts of shared genes. This integration strategy aimed to enhance the sensitivity of the analysis and mitigate false negatives associated with the limitations of each isolation protocol.

Functional enrichment analysis was performed on the merged dataset. First, a differential pathways analysis was performed using the Gene Set Variation Analysis (*GSVA*) R package [68]. Gene sets corresponding to Gene Ontology (GO) biological processes were obtained from the Molecular Signatures Database (MSigDB)□and used as the reference collection. Differentially expressed pathways (DEPs) were identified using the *limma* package□[69], applying an adjusted *P*-value threshold of <□0.05, with TP1 as the reference group. Additionally, we conducted an over-representation analysis (ORA) of GO terms with the *ClusterProfiler* R package [70, 71], *org.Hs.eg.db* [72], and *AnnotationDbi* [73]. These tools facilitated the identification of pathways significantly enriched among DEGs, providing insight into the biological processes modulated in response to the experimental conditions. For specific analyses, we generated a subset of neurobiological processes by systematically and manually curating the GO (biological processes) database, selecting entries that included neurobiologically related terms. This approach resulted in a total of 483 biological processes (**Table S1**).

We next compared the DEGs found in the merged dataset with those indexed in the Simons Foundation Autism Research Initiative (SFARI) database [74]. This database, developed for the ASD research community, compiles information on genetic variants associated with the etiology of ASD. Genes in the SFARI database are classified into four categories: *i*) syndromic (genes with mutations strongly linked to ASD and additional clinical features beyond core diagnostic criteria), *ii*) Level 1 (high-confidence ASD risk genes), *iii*) Level 2 (strong candidate genes), and *iv*) Level 3 (genes with moderate evidence based on previous studies). This comparison allowed us to assess the extent to which our findings align with the existing literature on ASD genetics.

### Co-expression modules analysis

To explore gene co-expression patterns potentially associated with musical stimuli in the buccal mucosa of ASD patients, we constructed a signed weighted correlation network using the Weighted Gene Co-expression Network Analysis (WGCNA) R package [75]. Gene expression data of the merged dataset were first normalized and corrected for inter-individual variability. To focus on the most informative signals, we retained the top 75% of genes with the highest variance across samples. A soft-thresholding power of 26 was selected based on scale-free topology criteria, yielding a model fitting index above 0.80 for both datasets (**Figure S1A**). We computed the Topological Overlap Matrix (TOM) and corresponding dissimilarities (1–TOM), followed by hierarchical clustering. Modules were defined using a minimum size of 30 genes and merged using a dendrogram cut height of 0.2 (**Figure S1B**). Each resulting module was assigned a unique color identifier. Module eigengenes were correlated with the TP1-TP2 to identify modules significantly associated with musical stimulation, based on gene significance (GS). Module membership (MM), reflecting intramodular connectivity, was used to identify hub genes (those with the highest connectivity). Each significant module was labeled according to its hub gene name.

To assess biological relevance, we conducted an over-representation analysis using the *compareCluster* function from the *clusterProfiler* R package [70], focusing on Gene Ontology (GO) biological process terms.

### Exploratory metagenomic analysis of salivary samples

To investigate potential shifts in bacterial composition associated with musical stimulation, we conducted an exploratory metagenomic analysis using reads that did not align to the human reference genome. These unmapped reads were extracted using the STAR aligner with the *--outReadsUnmapped Fastx* parameter. Read quality was assessed with *FastQC* and summarized using MultiQC, revealing consistently high base quality scores across all samples. Common metagenomic artifacts, such as per-base sequence content biases, sequence duplication, and adapter contamination, were detected. These issues were addressed using *Trimmomatic*, which efficiently removed adapter sequences and improved overall read quality. These unmapped reads were processed and classified against a bacterial genome database using the *CLARK* metagenomic classifier (v1.3.0.0) [76, 77], which provides a taxonomic profile based on discriminative *k*-mers (short DNA sequences of fixed length *k*) and assigns bacterial labels based on discriminative *k*-mer comparisons against the NCBI/RefSeq bacterial genome database. *CLARK* constructs an index from reference genomes by identifying unique *k*-mers that are specific to each genome. Shared *k*-mers among multiple genomes are excluded to ensure that only genome-specific sequences are used for classification. During the read assignment, *CLARK* quantifies the number of shared discriminative *k*-mers between each read and reference genomes, classifying each read to the genome with the highest match. The analysis was conducted in full mode, which utilizes the complete set of discriminative *k*-mers and outputs detailed metrics, including read counts per taxon and confidence scores (ranging from 0.5 to 1.0). This mode was selected to retain maximum resolution and accuracy in taxonomic assignments.

*DESeq2* package was used to normalize the data and detect differences in the abundance of microorganisms after the musical stimuli when compared to the baseline. We include the patient ID in the model as a random effect to account for patient-to-patient differences in the baseline using the *limma* package.

Graphical visualizations were generated in RStudio (v4.4.2) using the *ggplot2* and *ggwaffle* packages [78]. Visual outputs included bar plots of the most abundant bacterial genera and comparative plots of relative and absolute abundances before and after musical exposure.

## Results

### Quality assessment of raw data and comparative evaluation of the mapping from library preparation protocols

Initial quality control assessments revealed that the raw sequencing reads were of very high quality across all samples, with optimal values observed for most quality parameters (data not shown). However, two quality metrics were consistently flagged across all samples: “Per Base Sequence Content” and “Sequence Duplication Levels.” These warnings, though persistent, are not unusual in RNA-seq experiments and are not indicative of poor sample quality. In contrast, the presence of adapter contamination indicated a clear need for improvement. To address this, sequencing adapters and low-quality bases were removed using *Trimmomatic*. A comparative quality assessment between Human-Enriched and Poly-A RNA-seq libraries in both pre-processed and trimmed datasets revealed several differences across different sequencing metrics, reflecting intrinsic characteristics of the two library preparation strategies. The most remarkable difference between the two methods is related to the duplication rates. Human-Enriched library displayed higher levels of duplication, with a mean rate of 84.04% in the pre-processed data and 85.42% in post-trimming data (**Table S2**). In contrast, Poly-A libraries exhibited markedly lower duplication, with an average of 61.88% pre-trimming and a moderate increase to 67.89% after trimming. GC content also differed between the two library types. Human-Enriched library maintained a consistently higher GC percentage, averaging 52–53% across both pre-trimmed and trimmed data, but, in contrast, Poly-A library exhibited a lower GC content, ranging from 42.95% in the pre-processed data to 44.1% post-trimming (**Table S2**). Finally, a substantial reduction in failed reads was observed following trimming in both library types.

To evaluate mapping efficiency and alignment quality, the output log files from STAR were aggregated and analyzed using *MultiQC*. Overall, poor alignment performance was observed in both the Human-Enriched and Poly-A datasets, characterized by a high proportion of reads that failed to map to the human reference genome (**Figures 2A** and **2B**). This result was not unexpected, given the RNA source, saliva, which contains a substantial amount of microbial and other non-human genetic material. These unmapped reads were subsequently analyzed through a dedicated metagenomic pipeline to investigate the composition and modulation of the oral microbiome in response to musical stimulation. A comparative assessment of the two datasets revealed notable differences attributable to their respective library preparation methods. The Poly-A dataset generally yielded a higher total number of reads, both mapped and unmapped, as shown in **Figure 2C** (see also **Table S3**). However, despite this greater sequencing depth, a smaller proportion of reads in the Poly-A dataset successfully aligned to human genes compared to the Human-Enriched dataset (**Figure 2D**).

**Figure 2.**
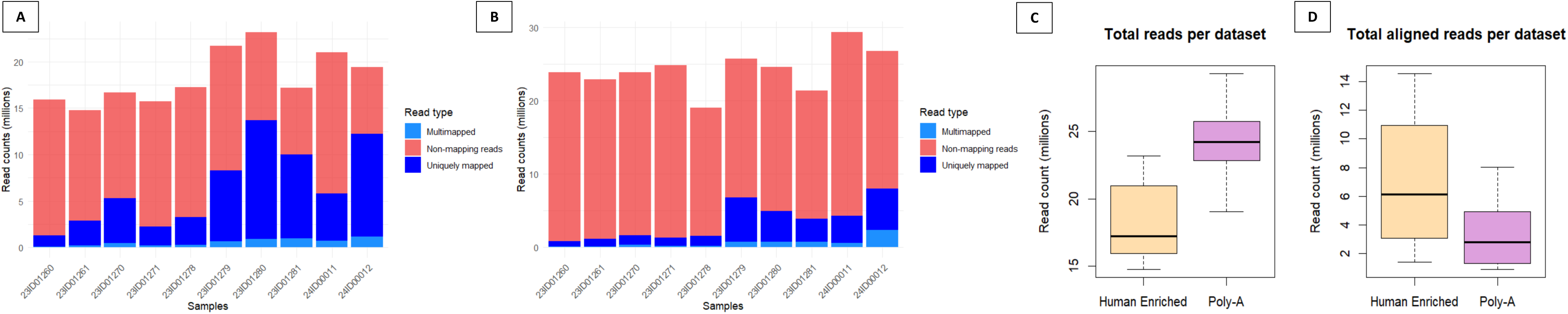
Sequencing quality and alignment metrics across datasets. Mapping statistics illustrating the alignment performance of the Human-Enriched dataset (A) and the Poly-A dataset (B) against the human genome (hg38). (C) Boxplots displaying the total number of reads per dataset in the Human-Enriched dataset and the Poly-A dataset. (D) Boxplots showing the total number of reads aligned to the human genome (hg38) in the Human-Enriched dataset and the Poly-A dataset.

Further analysis showed that the Poly-A dataset achieved broader transcriptomic coverage, detecting approximately 20,390 genes. However, this broader coverage came at the cost of lower average read counts per gene. In contrast, the Human-Enriched dataset, as expected, showed a more focused mapping pattern: while fewer genes were detected (approximately 17,730), the number of reads mapping to each gene was higher, indicating increased sequencing depth per gene.

### Differential gene expression analysis

A differential expression analysis was carried out to detect differentially expressed genes after the musical stimuli in the two salivary datasets separately. To improve statistical power and reduce noise, genes with fewer than 10 counts in at least five samples were filtered out. After applying this filtering criterion, 9,455 genes were retained in the Human-Enriched dataset, while only 6,791 genes were retained in the Poly-A dataset. The lower number of retained genes in the Poly-A dataset likely reflects its lower average read count per gene and sample.

Following statistical testing, the Human-Enriched dataset yielded 17 differentially expressed genes (DEGs) with adjusted *P*-values below 0.1, comprising 11 upregulated and 6 downregulated genes after musical stimulation. In contrast, the Poly-A dataset yielded only 5 DEGs, all of which were upregulated. When considering genes with nominal *P*-values < 0.05, the Human-Enriched and Poly-A datasets yielded 720 and 442 genes, respectively. Of these, 57 genes were shared between the two datasets. Among these common genes, 55 were protein-coding, one was a long non-coding RNA, and one was a transcribed unprocessed pseudogene.

To assess the concordance in expression trends between the two datasets, Log_2_FC for the 57 shared genes was compared. Pearson correlation analysis revealed a significant correlation between the Human-Enriched and Poly-A datasets (*R* = 0.77), indicating a moderate-to-high level of agreement between the two datasets (**Figure 3A**). This result suggests that despite technical differences and inherent biases between the two library preparation methods, both datasets successfully captured overlapping transcriptional responses to the musical stimulus.

**Figure 3.**
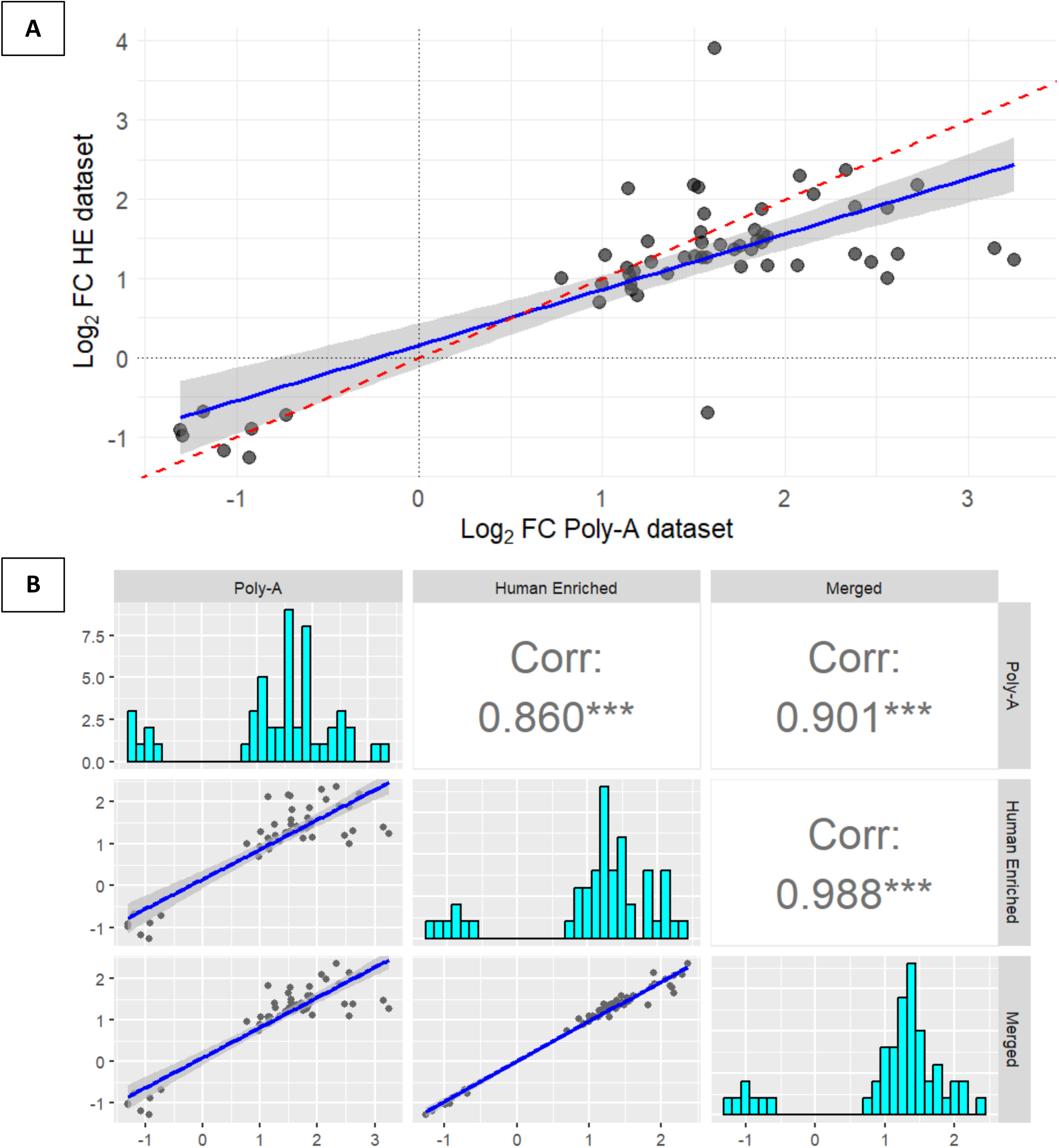
Correlation analyses between gene expression datasets to assess consistency in differential expression. (A) Correlation plot showing the Log_2_FC of 57 common genes (*P*-value < 0.05) between the Human-Enriched (HE) and Poly-A datasets (correlation = 0.77; 95% confidence interval). (B) Correlation plot of 50 genes (*P*-value < 0.05) shared between the merged dataset and the previously identified 57 common genes from the HE and Poly-A datasets.

### Integration through a merged dataset and cross-dataset validation of expression trends

To further optimize the analysis, a merged dataset was created by summing the read counts of shared genes across samples that were present in both datasets. This integration approach aimed to combine the broader gene coverage offered by the Poly-A dataset with the higher per-gene sequencing depth of the Human-Enriched dataset. Each dataset was filtered individually using the same criteria (genes with ≥10 counts in at least five samples). Afterward, counts from both datasets were merged by summing their values, followed by normalization and differential expression analysis using *DESeq2* (**Table S4**). To further confirm the consistency of expression patterns in the two datasets, genes with *P*-values below 0.05 in the merged dataset (*n* = 433) were intersected with the 57 genes previously found to be shared between the Human-Enriched and Poly-A datasets. A total of 50 genes were common to all three analyses. The Log_2_FC values for these genes were again compared across datasets. The Pearson correlation coefficient was 0.86 when comparing the Human-Enriched and Poly-A libraries, and exceeded 0.90 when each was compared individually with the merged dataset (Human-Enriched *vs*. Merged: 0.988; Poly-A *vs*. Merged: 0.901). All three comparisons were highly statistically significant (**Figure 3B**). These correlation values indicate strong consistency in expression patterns and further support the validity of the merging approach.

The merged dataset revealed three DEGs after musical stimuli when compared to baseline, with adjusted *P*-values below 0.1 (**Figure 4A**). Other genes previously associated to ASD were also significantly affected by the musical stimuli. Two genes, *HERC6* and *TSPAN5*, were upregulated, while *REM2* was downregulated (**Figure 4B**). Gene expression profiles of the 50 common genes could perfectly separate TP1 from TP2 in both the PCA (**Figure 4C**) and the heatmap analysis (**Figure 4D**)

**Figure 4.**
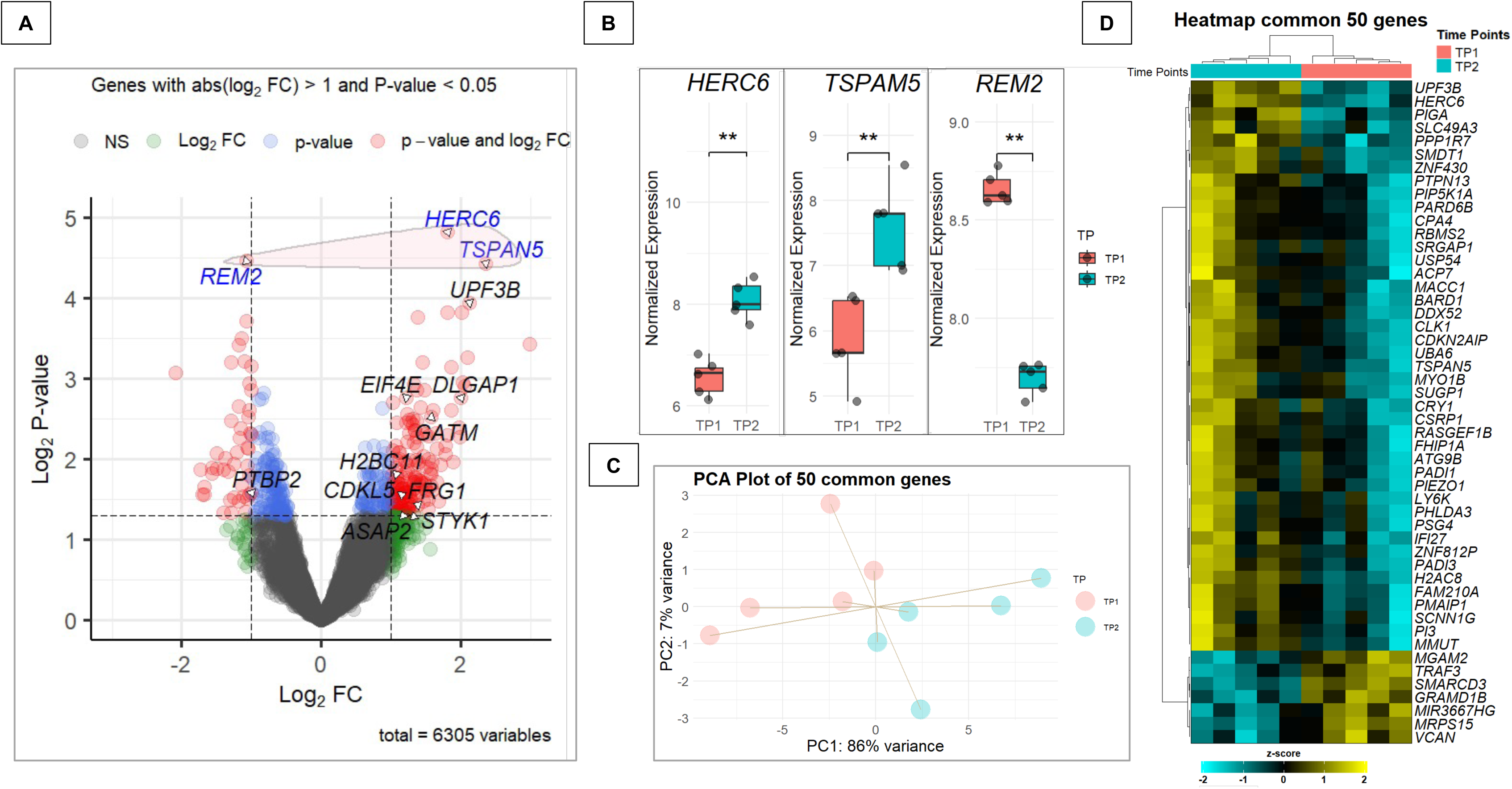
Analysis of DEGs and their relevance to autism-related pathways. (A) Volcano plot of the merged dataset, highlighting genes listed in the SFARI database and encircling in red the three genes with adjusted *P*-value < 0.1 that are not included in the SFARI database. (B) Boxplot showing the expression distribution of DEGs in the merged dataset (\*\**P*-value = 0.0079). (C) Principal Component Analysis (PCA) of the same 50 genes, illustrating sample distribution based on gene expression profiles. (D) Heatmap displaying the expression patterns of the 50 genes from the intersection between the genes (*P-value* < 0.05) in the merged dataset and the 57 genes previously found in common between the Human Enriched (HE) and the Poly-A datasets (*P*-value < 0.05).

### Functional Interpretation and Pathway Exploration

Although the number of statistically significant DEGs identified in this study was limited, the high consistency observed across multiple datasets and analytical approaches strongly supports the hypothesis that music exposure can elicit measurable transcriptional changes in the buccal mucosa of individuals with ASD. To further explore the potential biological relevance of these changes, we performed pathway-level analyses using Gene Set Variation Analysis (GSVA) and Over-Representation Analysis (ORA). These complementary methods allowed us to move beyond single-gene effects to identify broader biological pathways and molecular functions modulated by musical intervention.

The differential pathway analysis revealed 137 differentially regulated pathways (DRPs) in ASD patients (adjusted *P*-value < 0.05), of which 58% (*n* = 79) were upregulated and 42% (*n* = 58) were downregulated following music exposure (**Figure 5A**; **Table S5**). Among the most significantly affected pathways (|Log₂FC| > 0.5 and adjusted *P*-value < 0.01), we identified biological processes such as aorta development, actin filament severing, endoplasmic reticulum to cytosol transport, and skeletal muscle tissue regeneration. Notably, 13 of the 137 DRPs were associated with neurobiological functions (**Figure 5B; Table S5**).

**Figure 5.**
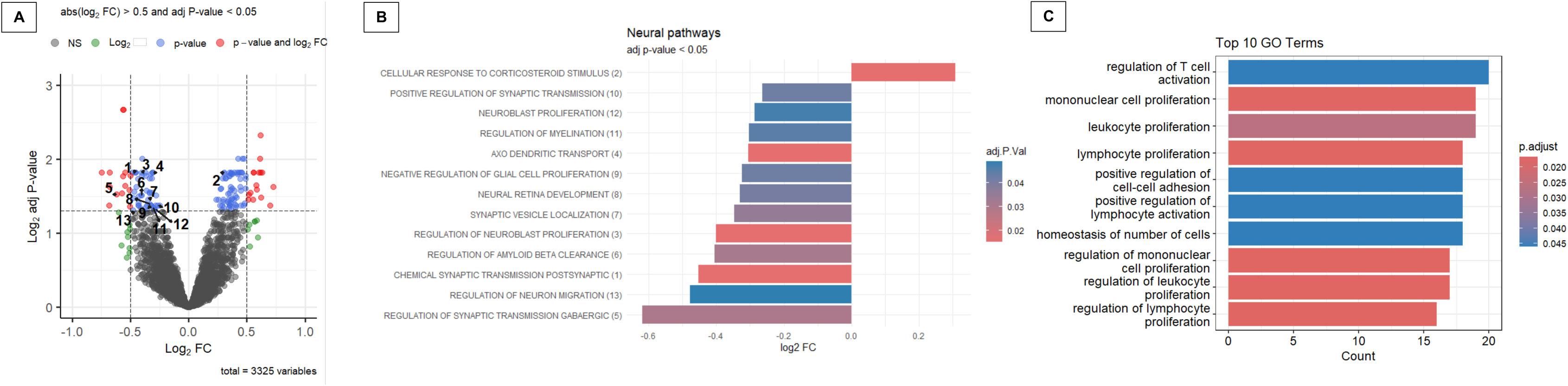
Comprehensive functional enrichment analysis of the merged dataset across multiple approaches. (A) Volcano plot of pathway alterations identified via Gene Set Variation Analysis (GSVA); The numbers in the plot correspond to GSVA-enriched neural functions. (B) Barplot of significantly GSVA-enriched pathways related specifically to neuronal biological functions (adjusted *P*-value < 0.05). (C) Barplot summarizing the top 10 significantly enriched pathways identified through Over-Representation Analysis (ORA).

Interestingly, in contrast to the global trend toward upregulation, neurobiological pathways showed a predominance of downregulation: 12 out of 13 exhibited negative Log₂FC values post-stimulation, suggesting reduced activity after music exposure. These included key processes such as chemical synaptic transmission, postsynaptic (Log₂FC = –0.45, adjusted *P*-value = 0.015), regulation of neuroblast proliferation (Log₂FC = –0.40, adjusted *P*-value = 0.015), and axo-dendritic transport (Log₂FC = –0.31, adjusted *P*-value = 0.018), pointing toward a potential dampening of excitatory synaptic signaling and neurodevelopmental activity. The only upregulated neurobiological pathway (Log₂FC = 0.31, adjusted *P*-value = 0.015) was the cellular response to corticosteroid stimulus. Additionally, regulation of GABAergic synaptic transmission was notably downregulated (Log₂FC = –0.62, adjusted *P*-value = 0.030). Other affected pathways involved in synaptic plasticity, neurodevelopment, and neurodegeneration, such as regulation of amyloid-beta clearance, synaptic vesicle localization, and neuron migration, were also significantly suppressed.

Functional analysis using the ORA approach identified 19 differentially regulated pathways (**Figure 5C; Table S6**), many of which were associated with immune-related processes. These pathways included those involved in immune cell proliferation and activation, migration and adhesion, as well as the maintenance of immune homeostasis.

### Co-expression modules analysis in response to musical stimulation

*WGCNA* analysis was conducted on a total of 4,667 genes from the merged dataset, after filtering out less variable genes. The analysis detected 28 modules of co-expressed genes, and among them, eight modules were significantly correlated with the musical stimuli. Multiple test correction was applied (FDR□<□0.05), resulting in three statistically significant correlated modules (|*R*| > 0.8); **Table S7**. Two of these, bisque4 (hub gene: *BCL10*) and saddlebrown (*UBE2D3*), exhibited strong positive correlation, while one module, plum1 (*AKNA*), showed a strong negative correlation with the music stimuli (**Figure 6A**). Other modules, such as darkolivegreen (*CDK2AP1*) and steelblue (*PIK3C2A*), also demonstrated notable correlation (*r* = – 0.75 and *R* = 0.76, respectively), although they did not retain statistical significance after correction (adjusted *P*-value = 0.065 for both). Equally, other additional modules, including ivory (*C1GALT1*), thistle1 (*UXS1*), and lightgreen (*FYN*) exhibited moderate correlations (*r* ranging from 0.64 to 0.67), but did not reach significance following adjustment (**Table S7**). Correlation between module membership (MM) and gene-level correlation with the musical stimuli of the three significant modules demonstrated a robust and statistically significant association, supporting the idea that core genes within these modules are functionally aligned with the musical response (**Figure 6B**). The most substantial association was observed in the *BCL10* module (*r* = 0.76; *P*-value = 6.3e-09). Heatmap analysis of gene expression patterns from genes included in each of the significantly associated modules evidenced their overall over-regulation (*BCL10* and *UBE2D3* modules) and under-regulation (*AKNA* module) after listening to music (**Figure 6C**). Eigengene values of individual samples also align with the module-trait correlation values and changes in expression patterns observed between TP1 and TP2 (**Figure 6B**).

**Figure 6.**
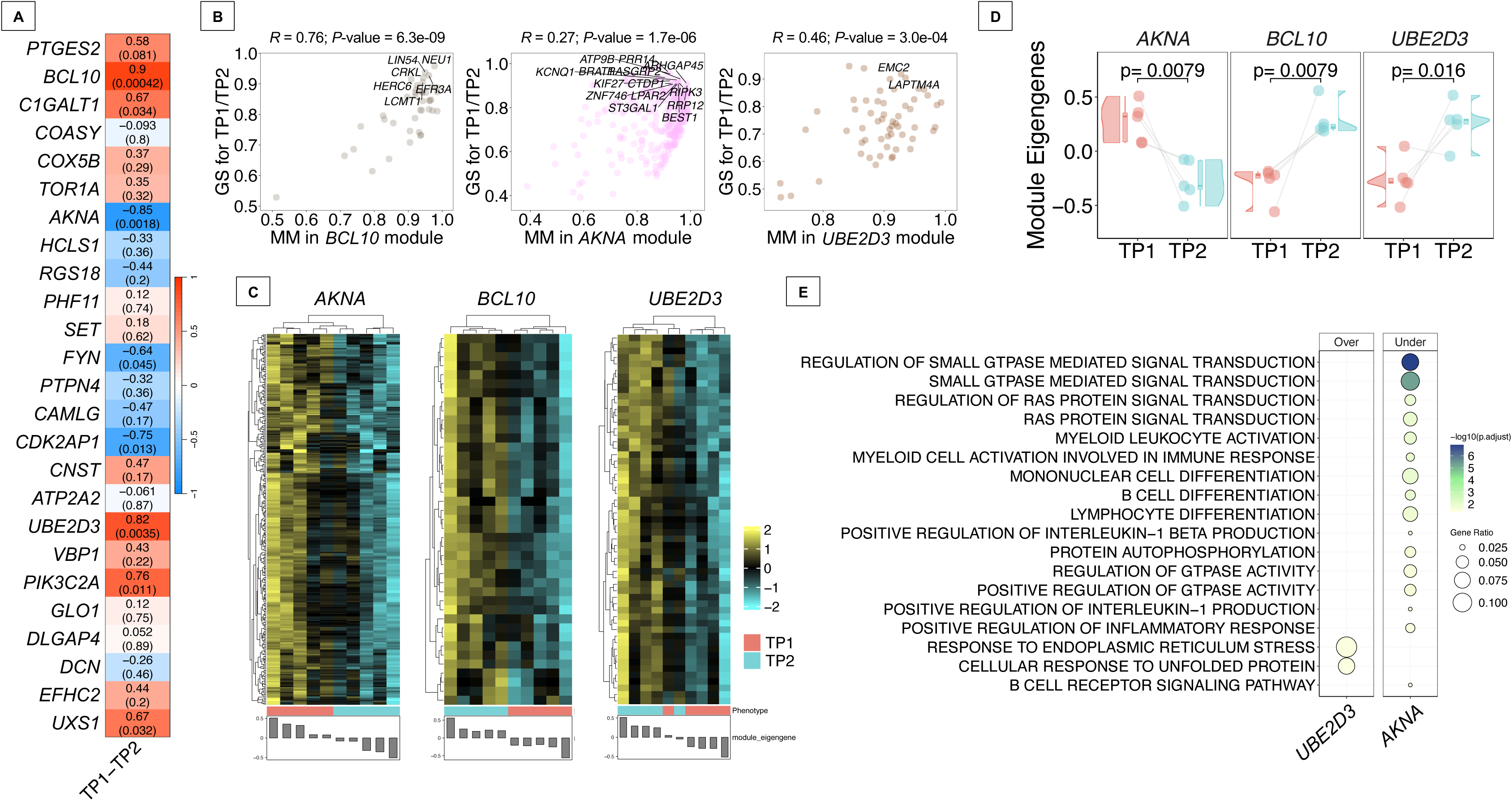
Co-expression analysis in ASD donors. (A) Heatmap of correlation values between modules of co-expressed genes detected and the musical stimuli (TP1-TP2). The upper value corresponds to the individual correlation value of each of the modules. *P*-values of these correlations are displayed in brackets. (B) Comparison between MM (module membership) and musical stimuli correlation of genes from the most significant module detected (FDR < 0.05). Only names from genes showing an MM > 0.9 and a correlation with music > 0.9 are displayed. (C) Heatmap of gene expression profiles from the most significant module detected (FDR < 0.05). (D) Raincloud plots of differences in individual samples’ eigengenes between TP1 and TP2 from modules showing statistical significance (FDR < 0.05). (C) Biological processes detected from most statistically significant modules (FDR < 0.05). Modules with significant pathways are named as the hub gene names.

Functional enrichment analysis identified different processes associated with the significantly correlated modules *AKNA*, *FYN*, *UBE2D3* and *UXS1* (**Table S8**). Notably, the *AKNA* and *UBE2D3* modules were among the three modules significantly correlated with the musical stimuli after false discovery rate (FDR) correction (FDR□<□0.05); **Figure 6C**. The *AKNA* module, which was downregulated in response to the stimuli, was primarily enriched for processes involving GTPase-mediated signal transduction, including RAS protein signaling pathways. Additionally, this module was also associated with immune-related functions such as myeloid cell differentiation and activation, B-cell differentiation and signaling, and the regulation of inflammatory responses, including interleukin-1 (IL-1) production (**Figure 6C**). On the other hand, only two enriched pathways were identified among the genes included in the upregulated *UBE2D3* module, namely ‘response to endoplasmic reticulum stress’ and ‘cellular response to unfolded protein’.

### ASD-related genes *vs*. genes altered by musical stimuli

To explore the potential relevance of musical stimulation to autism-related molecular mechanisms, we compared the gene expression data from our merged dataset with the SFARI gene database. Of the 6,305 genes identified, 519 matched entries in the SFARI database. Among these, ten genes showed significant differential expression (*P*-value < 0.05 and |Log_2_FC| > 1): nine were upregulated and one was downregulated. Notably, genes such as *EIF4E*, *DLGAP1*, and *CDKL5* were among the upregulated group, while *PTBP2* was significantly downregulated. Several of these genes have strong or suggestive associations with autism, according to their SFARI scores (e.g., *EIF4E* – Score 2; *CDKL5* – Score 1S). The identified genes are implicated in various processes, including synaptic signaling, protein translation, and RNA processing, suggesting that music stimulation may influence molecular pathways commonly affected in ASD (**Table S9**).

### Exploratory analysis of salivary microbiota following musical stimulation

We also investigated whether musical stimulation induces detectable changes in the salivary microbiota by performing metagenomic inference on reads that failed to align to the human genome. As expected, a substantial proportion of these unmapped reads could not be matched to any bacterial reference genome (**Figure S2**). This result is consistent with the complex and heterogeneous composition of saliva, which contains not only bacteria but also fungi, viruses, dietary RNA fragments, and environmental contaminants that may not be represented in the reference database used.

Applying a filtering criterion consistent with the human differential gene expression (DGE) analysis (retaining only species with ≥10 reads in at least five samples), the number of species was reduced from 3,210 (Human-Enriched dataset, HE) and 3,345 (Poly-A dataset) to 1,340 and 1,767 species, respectively. A total of 1,273 species were shared between both datasets.

Differential abundance analysis (DAA) was conducted using *DESeq2* in the two datasets separately. No bacterial species passed the adjusted significance threshold (adjusted *P*-value < 0.05) in either dataset. However, 52 species in the Human-Enriched dataset and 62 in the Poly-A dataset showed nominal significance (*P*-value < 0.05), with 15 species overlapping between the two datasets: *Austwickia chelonae, Propionibacterium freudenreichii, Capnocytophaga sp. Oral taxon 864, Brachybacterium vulturis, Kingella denitrificans, Laribacter hongkongensis, Kingella kingae, Sulfuriferula plumbiphila, Usitatibacter palustris, Ephemeroptericola cinctiostellae, Helicobacter winghamensis, Hathewaya histolytica, Bacteroides faecium, Arcanobacterium buesumense*, and *Acidipropionibacterium acidipropionici*. A Pearson correlation analysis of the Log_2_FC for these 15 shared species revealed a high concordance (*r* = 0.93), indicating strong agreement between the independent analyses (**Figure 7A**).

**Figure 7.**
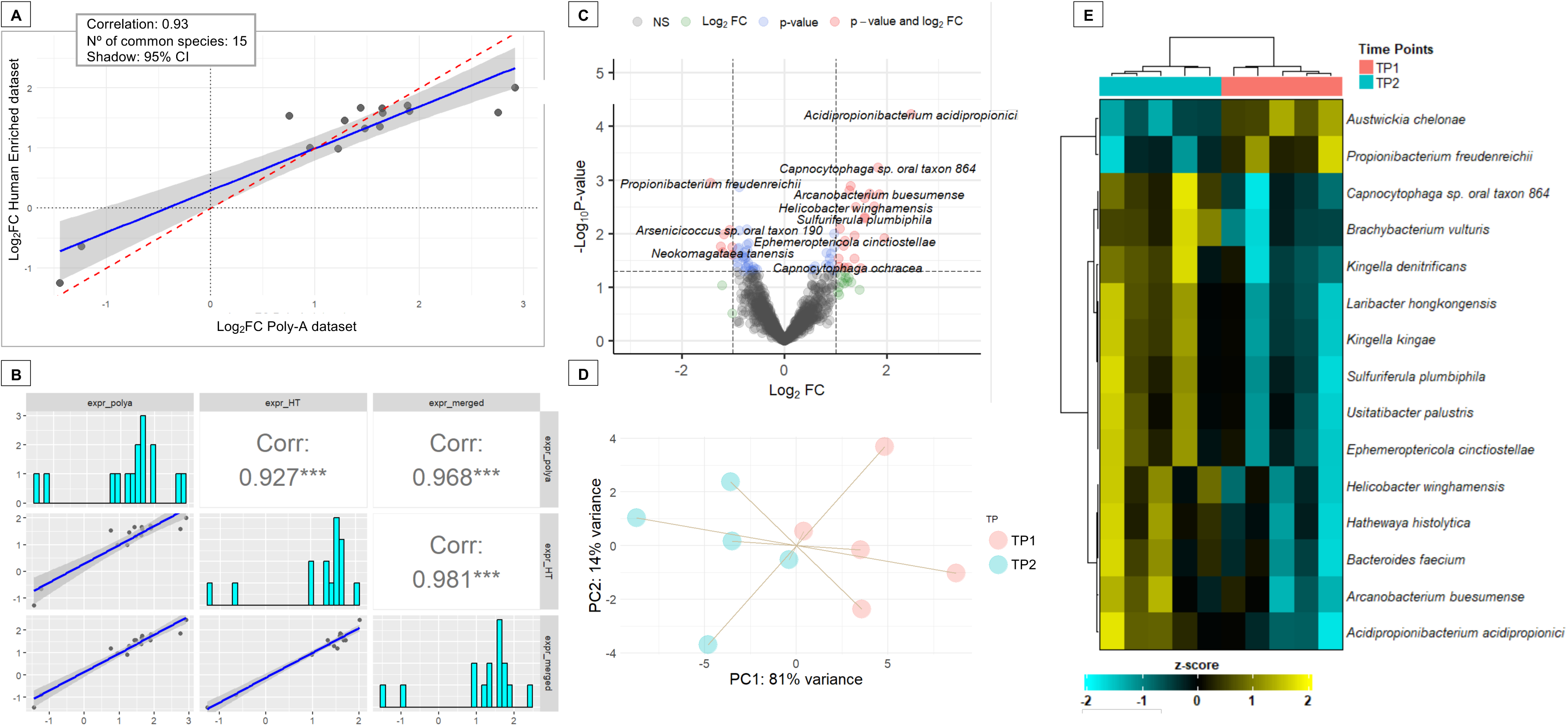
Differential abundance analysis of salivary microbial species pre- and post-musical stimulation in ASD subjects. (A) Scatter plot showing the Pearson correlation of Log_2_FC values for 15 microbial species with differential abundance in both the Human-Enriched (HE) and Poly-A datasets. (B) Correlation plot of the 15 species from the intersection from the merged dataset and previously found species in common between the Human-Enriched and the Poly-A datasets, which were 15. (C) Volcano plot of the merged dataset displaying species with significant differential abundance. (D) PCA plot obtained from the same 15 species in the merged dataset (E) Heatmap of the relative abundances of the 15 shared species between HE and Poly-A datasets.

Subsequently, a merged dataset was generated by aggregating read counts across datasets for the 1,273 shared species. This merged dataset showed also a high correlation in Log_2_FC values with the Poly-A dataset and Human-Enriched datasets for the 15 shared species between the two libraries (**Figure 7B**). Differential abundance analysis (DAA) identified 68 species with significantly different abundances at TP2 compared to TP1 (*P*-value < 0.05; **Table S10**; **Figure 7C**). Notably, all 15 species previously found in common between the Human-Enriched and Poly-A datasets were also detected in the merged dataset. Changes in the abundance of these 15 species effectively distinguished the pre-stimulation and post-stimulation states in ASD subjects, as demonstrated by both PCA (**Figure 7D**) and heatmap analysis (**Figure 7E**). Unlike the separate analyses, this integrated approach identified one additional differentially abundant species, *Acidipropionibacterium acidipropionici*, with an adjusted *P*-value of 0.076 and a Log_2_FC of 2.5 (**Table S10**).

## Discussion

This study explored the transcriptional effects of music exposure in the buccal mucosa of individuals with ASD using two distinct RNA sequencing datasets prepared *via* Poly-A selection and Human-Enriched library protocols. Human-Enriched and Poly-A RNA-seq libraries exhibited distinct sequencing characteristics that align with their differing biological targets and preparation protocols. Human-Enriched library showed higher duplication and GC content compared to Poly-A library preparation, and although it detected fewer genes, it achieved higher read counts per gene, reflecting greater sequencing depth for targeted transcripts. This is expected, as Poly-A generally captures more genes than Human-Enriched because it uses poly-A selection to isolate all polyadenylated transcripts, allowing for an unbiased, transcriptome-wide analysis of coding genes. Thus, while Poly-A enriches for the majority of protein-coding mRNAs expressed in the samples without restriction to predefined regions (enabling detection of a broad range of transcripts across the genome), Human-Enriched relies on hybrid capture with a targeted panel (e.g., the Exome Panel), limiting analysis to a specific subset of genes or regions. Although each dataset individually revealed a poor mapping performance against the human genome and a modest number of DEGs, the integration of both datasets enabled the identification of a more consistent transcriptional signature, with enhanced robustness and biological interpretability. The moderate-to-high correlation (*r* = 0.77) in gene expression trends between the Poly-A and Human-Enriched datasets, and the high consistency (*r* > 0.90) in the merged dataset, reinforce the validity of the transcriptional changes observed. Although the number of DEGs detected at stringent thresholds was limited, likely due to sample size constraints and biological variability inherent in human studies, the robustness of these shared signals underscores the value of integrating different RNA-seq strategies.

Three genes, *HERC6*, *TSPAN5*, and *REM2*, emerged as significant from the merged Poly-A selection and Human-Enriched datasets, suggesting they may play important roles in the brain’s response to musical stimuli in individuals with ASD. The upregulation of *HERC6* may point to a role for immune-related pathways in mediating the effects of music exposure. *HERC6* encodes an E3 ubiquitin-protein ligase and is a member of the small HERC family, which, unlike its larger counterparts (*HERC1*, *HERC2*), is thought to have evolved independently. While in many species *HERC6* participates in type I interferon responses by facilitating ISGylation through the ubiquitin-like protein ISG15, this function appears to be lost in humans, where ISGylation activity is instead mediated by *HERC5* [79]. Despite this loss, emerging evidence suggests that *HERC6* may still play roles in antiviral defense and immune regulation through alternative, non-ISGylation-based mechanisms [80]. The modulation of such immune-related genes is particularly relevant given accumulating evidence that immune dysregulation is a common feature in ASD. Music, by influencing stress levels and immune signaling pathways, could indirectly affect the local expression of genes such as *HERC6*, potentially contributing to homeostatic recalibration in neuroimmune interactions. Perhaps more directly tied to neurodevelopmental processes, the upregulation of *TSPAN5* presents a compelling finding. *TSPAN5* encodes a glycoprotein of the tetraspanin family, which is involved in a broad array of cellular functions including adhesion, migration, and proliferation. Of particular importance is its role in the central nervous system, where it is highly expressed in the forebrain and cerebellum. Recent studies have demonstrated that *TSPAN5* facilitates the maturation of dendritic spines by promoting the synaptic clustering of neuroligin-1 (NLG-1) [81], a protein that has been closely linked to ASD-related social behavior deficits [82]. The observed upregulation of *TSPAN5* in the buccal mucosa of ASD patients following musical intervention may thus reflect a shift toward synaptic stabilization and maturation occurring in the brain. This aligns with reports that music-based therapies can improve cognitive and social functioning in individuals with ASD, potentially through activity-dependent plasticity and enhanced synaptic integration. Conversely, *REM2* was significantly downregulated after music exposure. *REM2* belongs to the RGK subfamily of Ras-like GTPases and plays a critical role in neuronal differentiation and synaptic development. Experimental studies have shown that *REM2* knockdown reduces dendritic spine density and maturity, factors often associated with atypical connectivity in ASD. However, intriguingly, inhibition of *REM2* has also been associated with increased dendritic branching and arborization, resulting in more structurally complex neurons [83]. This dualistic effect complicates the interpretation of its downregulation, yet suggests that reduced *REM2* expression may contribute to neuroplastic adaptations rather than dysfunction per se. In the context of this study, the simultaneous upregulation of *TSPAN5* may offset potential drawbacks of *REM2* suppression, together fostering a structural environment conducive to improved neural integration and function.

The concurrent modulation of *HERC6*, *TSPAN5*, and *REM2* supports the hypothesis that music exposure can elicit biologically meaningful changes at the molecular level in the salivary transcriptome of individuals with ASD. These changes appear to span multiple functional domains, including immune regulation, synaptic maturation, and structural remodeling, all of which are relevant to the ASD phenotype. While the directionality and interplay between these pathways remain to be fully elucidated, our findings resonate with a growing body of literature suggesting that music can serve as a multisensory and emotionally salient stimulus capable of driving neuroplasticity.

Numerous studies have explored the relationship between ASD and alterations in the synaptic excitation-inhibition (E/I) ratio. Some have reported an increased E/I ratio in individuals with ASD [84], while others, particularly in the context of Down syndrome, a condition sharing certain phenotypic and cognitive features with ASD, have suggested that heightened inhibition may be the primary contributor to learning deficits [85]. Collectively, these findings seem to indicate that homeostatic regulation of neural activity is often disrupted in ASD. Given the heterogeneity of the disorder, it is plausible that both hyperexcitation and hyperinhibition could give rise to distinct ASD subtypes or manifestations. Pathway analysis revealed that musical stimulation led to a coordinated downregulation of neurodevelopmental and synaptic pathways in the buccal mucosa of ASD patients, contrasting with the global upregulation pattern observed across all GSVA-enriched pathways. In light of our GSVA results and the supporting literature, we hypothesize that the observed downregulation of most neural DRPs following musical stimuli may represent a homeostatic response aimed at modulating excessive synaptic excitation, a characteristic frequently observed in ASD. This response could reflect a reduction in the E/I ratio, underscoring the potential of music, especially when presented in a relaxing and familiar context, as a modulator of neural excitability. The cellular response to corticosteroid stimulus was the only significantly upregulated pathway, which may reflect a stress-related or neuroendocrine response to the stimulus. In contrast, our ORA revealed significant enrichment in biological pathways associated with immune processes, particularly those involving the proliferation and differentiation of immune cells such as leukocytes, lymphocytes, and mononuclear cells. Immune dysregulation is a well-established co-occurring condition in ASD, with several studies reporting elevated levels of TNF-α, IL-6, granulocyte colony-stimulating factor (G-CSF), IFN-γ, and IL-8 in the brain, along with a heightened pro-inflammatory Th1 cytokine profile [86–88]. The observed modulation of pathways related to immune cell differentiation following music exposure may suggest an attenuation of such pro-inflammatory states, supporting previous findings that music can exert immunoregulatory effects and influence gene expression related to immune and stress responses.

While further mechanistic studies are warranted, our GSVA and ORA results support the notion that external sensory stimuli such as music may trigger compensatory transcriptional programs in individuals with ASD, contributing to the regulation of both neural excitability and immune function. By examining the DEGs stimulated by music in ASD patients and cross-referencing them with the SFARI gene database, we additionally found that music stimulation appears to modulate a network of genes involved in synaptic structure, intracellular signaling, and RNA metabolism, pathways commonly disrupted in ASD. Notably, several upregulated genes, including *DLGAP1* and *CDKL5*, play critical roles in synaptic scaffolding and neuronal morphogenesis [89, 90]. *DLGAP1* encodes a postsynaptic protein localized at glutamatergic synapses, while *CDKL5* is essential for the stability of excitatory synapses [90]. The altered expression of these genes suggests a potential link between musical stimulation and enhanced synaptic plasticity in ASD-related contexts. In addition, genes such as *EIF4E*, *STYK1*, and *CDKL5* are components of the PI3K/Akt/mTOR signaling pathway, a key regulator of synaptic protein synthesis and neural development. Their upregulation supports the hypothesis that music may influence translational control mechanisms implicated in ASD pathophysiology. Importantly, we also observed the downregulation of *PTBP2*, a central regulator of neuronal exon splicing and RNA stability. Along with other RNA-binding genes such as *UPF3B* and *FRG1*, this finding indicates that music may exert broader effects on RNA processing and transcript diversity [91, 92]. Taken together, these transcriptional changes suggest that music stimulation can reprogram gene expression patterns across interconnected molecular pathways, potentially contributing to neuroplastic and compensatory mechanisms relevant to ASD.

We identified different co-expression modules whose activity patterns were significantly correlated with exposure to musical stimuli in ASD donors, suggesting that musical stimuli may engage specific regulatory networks in the buccal mucosa, potentially reflecting broader systemic or brain-related transcriptional responses. From the three main correlated modules (adjusted P-value < 0.05), *BCL10* and *UBE2D3* showed a global upregulation pattern in response to music stimulation, while the *AKNA* module demonstrated an overall downregulation. Notably, hub gene *AKNA* has already been previously reported as a novel ASD risk gene, carrying *de novo* variants probably related to ASD [93]. *AKNA* module was functionally enriched for pathways related to GTPase-mediated signal transduction, with a particular emphasis on Ras protein signaling, suggesting that musical stimulation may dysregulated specific intracellular signaling cascades related to these small GTPases in the buccal mucosa of ASD subjects. The Rho family of GTPases (members of the larger Ras superfamily) are small signaling proteins that play pivotal roles in neural development, including axon guidance, dendritic spine formation, and synaptic plasticity processes (REF). Importantly, dysregulation of Rho GTPase signaling has been increasingly recognized as a contributing factor in various neuropathological events, including ASD [94, 95]. Multiple lines of evidence have highlighted aberrant activity of Rho family members, such as Rac1, in ASD-related phenotypes both in human studies and animal models [96]. In addition, this module was also enriched for genes involved in immune-related processes, suggesting an immunomodulatory effect of the musical stimuli at the local level, particularly through the suppression of different cells involved in immune responses. The downregulation of inflammatory and IL-1 pathways could be of particular interest given the role of neuroinflammation in the pathophysiology of ASD [97–99]. Module analysis also revealed a selective activation of stress-response mechanisms (*via* the *UBE2D3* module). The accumulation of unfolded proteins in the endoplasmic reticulum (ER) triggers ER stress, specifically activating a protective unfolded protein response in the ER. Several studies have identified ER stress as a significant contributor to the pathophysiology of ASD [100]. ER stress has also been shown to impair neurite outgrowth, alter synaptic protein expression, and disrupt neuronal differentiation, all of which are relevant to ASD neurodevelopmental deficits [101]. In addition, upregulation of ER stress-related genes has been reported in the brains of ASD individuals [101, 102]. Our results suggest that music exposure may enhance ER stress response mechanisms in buccal cells, potentially regulated by genes within *UBE2D3* module, which could help to compensate for the impact of stress pathways implicated in ASD.

While the metagenomic analysis conducted in this study was exploratory and limited in scope, it yielded several notable observations. The large fraction of sequencing reads that failed to align with bacterial reference genomes highlights the inherent complexity of salivary metagenomic samples. Saliva is a biologically diverse medium that includes bacteria, fungi, viruses, dietary residues, and environmental RNA fragments, many of which are absent from current bacterial reference databases. This highlights the necessity for future metagenomic studies to employ more comprehensive taxonomic frameworks, ideally incorporating multi-kingdom reference sequences. Furthermore, improved annotation pipelines capable of integrating fungal, viral, and non-coding elements would enhance biological interpretation. Dysbiosis of the oral microbiome has previously been linked to ASD, potentially contributing to neurodevelopmental alterations via the gut–brain axis or immune modulation [103, 104]. A subset of taxa showed consistent nominal differences between pre- and post-stimulation samples across both datasets. In particular, 15 species demonstrated shared differential abundance patterns with high correlation in Log_2_FC (r = 0.93), suggesting reproducibility across sequencing approaches (Human-Enriched *vs*. Poly-A datasets). Notably, the merged dataset identified *Acidipropionibacterium acidipropionici* as a potentially modulated species following musical stimulation (adjusted *P*-value = 0.076). This acid-producing bacterium is known for its role in dairy fermentation [105], and closely related propionibacteria have been previously identified as part of the oral microbiota [106]. We also identified *Propionibacterium freudenreichii*, a close relative involved in propionic acid production, among our top 15 candidate species. There is substantial evidence implicating propionibacteria in neurological effects via propionic acid production, an important short-chain fatty acid. Elevated propionic acid levels in the gut have been associated with ASD-like behaviors and neuroinflammation in animal models [107]. According to Li et al. [108], 3-(3-hydroxyphenyl)-3-hydroxypropionic acid induces autism symptoms by emptying and depleting catecholamines in the brain. El-Ansary et al. [109, 110] reported that the neurotoxicity of propionic acid could play a central role in the etiology of autistic biochemical features. This finding raises the intriguing possibility that auditory stimulation may exert an “echo effect” on the composition of the oral microbiome. These results invite further inquiry into the interplay between sensory stimuli and microbial communities. It remains unclear whether the observed microbial shifts are directly mediated by neural or physiological responses to music, or indirectly through changes in salivary secretion, flow rate, or composition. Future studies with larger cohorts, increased sequencing depth, and broader reference annotations, including non-bacterial taxa, are needed to validate and expand on these preliminary findings. Further research should explore the potential of targeting the microbiota–gut–brain axis through musical stimuli as a therapeutic strategy in ASD, in parallel with approaches such as probiotic supplementation [108, 111], or more broadly, interventions that manipulate the enteric microbiome to alleviate autism symptoms [112].

One of the most significant challenges encountered in the present study was the use of saliva as the biological matrix for transcriptomic analysis. While saliva sampling offers clear advantages, being non-invasive, stress-free, and highly accessible, particularly in sensitive populations such as individuals with ASD, it also presents substantial technical hurdles. Saliva contains a heterogeneous mix of host epithelial cells, immune cells, and a rich microbial community, which can result in low yields of high-quality human RNA and high proportions of unmapped or non-human reads. These factors complicate both the quality control and the interpretation of gene expression data. Despite these challenges, this study demonstrates that with careful preprocessing, rigorous quality filtering, and thoughtful experimental design, it is possible to extract biologically meaningful transcriptomic signals from saliva. This opens the door for future research into peripheral biomarkers of brain function and neuroplasticity in contexts where blood collection is difficult or ethically sensitive, such as pediatric or neurodiverse populations. To further advance the utility of saliva transcriptomics, future studies should aim to improve RNA stabilization protocols, integrate host-microbiome transcriptome analyses, and consider single-cell or spatial transcriptomics to better resolve cell-type-specific expression patterns. Combining saliva-based transcriptomics with behavioral, neuroimaging, or immune profiling could yield a more comprehensive understanding of how systemic biological pathways contribute to neurodevelopmental conditions like ASD.

This study is not without limitations. The sample size, while reasonable for a pilot analysis, limits the power to detect subtle effects. The exclusion of outlier samples was necessary to reduce noise but may have inadvertently excluded biologically informative variance. Additionally, the focus on peripheral tissue (saliva) restricts direct extrapolation to brain-specific mechanisms, although saliva has increasingly been recognized as a viable proxy for systemic and neuroimmune interactions. Future studies should build upon these findings by increasing cohort sizes, incorporating longitudinal designs, and using complementary modalities such as neuroimaging and behavioral assessments. Furthermore, investigating cell-type-specific transcriptomic responses or incorporating single-cell RNA-seq could help disentangle the contributions of diverse cell populations and improve the resolution of observed effects.

In conclusion, this study offers preliminary yet compelling evidence that music exposure can modulate the expression of genes involved in synaptic plasticity, neuronal architecture, and immune signaling in individuals with ASD. Notably, the upregulation of *TSPAN5* and *HERC6*, alongside the downregulation of *REM2*, suggests a transcriptional profile that may favor enhanced neural adaptability and integration. These molecular findings enrich our understanding of the biological mechanisms through which music may exert therapeutic effects and point to promising avenues for developing targeted, non-invasive interventions in neurodevelopmental conditions. Furthermore, given the frequent reports of oral microbiota dysbiosis in ASD, the observed trends in microbial composition following musical stimulation, though exploratory, raise important questions about the broader systemic effects of sensory inputs. Understanding how music may influence the oral microbiome could open new perspectives on modulating host–microbe interactions through behavioral or environmental stimuli. Nevertheless, these initial findings should be interpreted with caution, and future research with larger cohorts and expanded metagenomic approaches will be essential to confirm and contextualize these effects.

## Acknowledgements

The authors would like to express their appreciation to the study investigators of the Sensogenomics network (sensogenomics.com; Sensogenomics Working Group [see Annex]), as well as the nursery and laboratory service at the Hospital Clínico Universitario de Santiago de Compostela, for their invaluable dedication and support. This research project was made possible through the access granted by the Galician Supercomputing Center (CESGA) to its supercomputing infrastructure. The supercomputer FinisTerrae III and its permanent data storage system have been funded by the Spanish Ministry of Science and Innovation, the Galician Government, and the European Regional Development Fund (ERDF). This work was supported by: *i*) GAIN IN607B 2020/08 and IN607A 2023/02, and EUTERPE_adn (Programa de Cooperación Interreg-VI POCTEP; Ref. 0313_EUTERPE_ADN_1_E) (to A.S.), IIN607A2021/05 (to F.M.-T.) and IN677D 2024/06 (to A.G.-C.), and *ii*) Consorcio Centro de Investigación Biomédica en Red de Enfermedades Respiratorias (CB21/06/00103; to A.S. and F.M.-T.). AG-C is supported by the Miguel Servet contract (CP23/00080), funded by the Instituto de Salud Carlos III (ISCIII) and co-funded by the European Union. The funders were not involved in the study design, collection, analysis, interpretation of data, the writing of this article, or the decision to submit it for publication.

## Supplementary Data

**Figure S1**. (A) Plots showing correlation between soft-thresholding powers and both the scale-free fit index and the mean connectivity (upper). (B) Dendrogram of genes and co-expression modules detected, represented by different colors.

**Figure S2**. Pie charts representing the percentage of bacterial reads and unknown reads in the Human-Enriched dataset (A) and in the Poly-A dataset (B)

**Table S1.** Neurobiologically related terms retrieved from the Gene Ontology database (GO).

**Table S2**. Quality control analysis of the reads from the Human-Enriched (HE) and Poly-A dataset, before and after trimming. For each sample (forward read R1 and reverse read R2 files), the table reports: percentage of duplicate reads (% Dups), average GC content (% GC), average read length (Read Length), percentage of failed reads (% Failed), and the total number of sequences in millions (M-seqs).

**Table S3**. Mapping statistics from STAR aligner of the reads from the Human-Enriched dataset and the Poly-A dataset. For each sample, the table reports the total number of sequenced reads (Total.reads), the number and percentage of uniquely aligned reads (Uniq.aligned, Uniq aligned %), and the number and percentage of multimapped reads (Multimapped, Multimapped %) after alignment to the human genome. The tables also include the average length of mapped reads (Avg. mapped len), the mismatch rate (Mismatch rate), deletion rate, and average deletion length (Del rate, Del len), and insertion rate and average insertion length (Ins rate, Ins len).

**Table S4**. Results of differential expression analysis from the merged dataset. The table includes the average expression level of a gene across all samples after accounting for library size differences (baseMean), the logarithm (base 2) of the fold change in gene expression after the musical stimuli (log2FoldChange), the standard error associated with the log2FoldChange (lfcSE), the Wald statistic, a measure of the difference in gene expression normalized by its standard error (stat), the unadjusted p-value associated with the Wald statistic (*P*-value), and the adjusted *P*-value, corrected for multiple testing using the Benjamini-Hochberg method.

**Table S5**. Significant pathways (adjusted *P*-value < 0.05) found in the Gene Set Variation Analysis (GSVA). The table reports, for each pathway, the log2 fold change in enrichment between conditions (logFC), the average GSVA enrichment score across all samples (AveExpr), the t-statistic from the linear model used to test differential enrichment (t), the P-value (*P*-value) and the adjusted P-value, differential enrichment based on a Bayesian mode (B). Lastly, the neuro column indicates the pathways related to neural processes, and the ID column shows the ID associated with each neural pathway.

**Table S6**. Significant pathways (adjusted P-value < 0.05) obtained from the Over Representation Analysis (ORA). For each pathway, the table reports the ID of the pathway and a short description, the number of associated genes (GeneRatio and Count), and how often the term appears in the background gene set (BgRatio). RichFactor and FoldEnrichment indicate how enriched the term is compared to what would be expected by chance. zScore measures the strength and direction of enrichment, while *P*-value, adjusted *P*-value, and Q-value show the statistical significance (corrected for multiple testing). Gene ID lists the genes associated with the term.

**Table S7.** Correlation between co-expression modules detected and the musical stimuli (TP1-TP2). P-values and adjusted P-values of the correlation were also calculated

**Table S8**. Clustered over-representation analysis from significantly correlated modules was detected. For each pathway, the table reports the ID of the pathway and a short description, the number of associated genes (GeneRatio and Count), and how often the term appears in the background gene set (BgRatio). *P*-value, adjusted *P*-value, and Q-value show the statistical significance (corrected for multiple testing). Gene ID lists the genes associated with the term.

**Table S9.** Differentially expressed genes (*P*-value < 0.05 &|Log_2_FC| > 1) in the merged dataset that found a match in the SFARI database, which contains genes associated with ASD. The table includes each gene’s average expression (baseMean), Log_2_ fold change (Log_2_FC), standard error (lfcSE), test statistic (stat), raw *P*-value, and adjusted *P*-value.

**Table S10**. Differential abundance analysis (DAA) results from the merged dataset. The table includes each gene’s average expression (baseMean), Log_2_ fold change (Log_2_FC), standard error (lfcSE), test statistic (stat), raw *P*-value and adjusted *P*-value.

## Sensogenomics Working Group

Antonio Salas Ellacuriaga – PI; Federico Martinón-Torres – PI; Laura Navarro Ramón – Coordinator

### GenPoB/GenVip – Instituto de Investigación Sanitaria (IDIS) (alphabetical order)

Alba Camino Mera, Albert Padín Villar, Alberto Gómez Carballa, Alejandro Pérez López, Alicia Carballal Fernández, Ana Cotovad Bellas, Ana Isabel Dacosta Urbieta, Narmeen Mallah, Ana María Pastoriza Mourelle, Ana María Senín Ferreiro, Andrés Muy Pérez, Antía Rivas Oural, Antonio Justicia Grande, Antonio Piñeiro García, Anxela Cristina Delgado García, Belén Mosquera Pérez, Blanca Díaz Esteban, Carlos Durán Suárez, Carmen Curros Novo, Carmen Gómez Vieites, Carmen Rodríguez-Tenreiro Sánchez, Celia Varela Pájaro, Claudia Navarro Gonzalo, Cristina Serén Trasorras, Cristina Talavero González, Einés Monteagudo Vilavedra, Estefanía Rey Campos, Esther Montero Campos, Fernando Álvez González, Fernando Caamaño Viñas, Francisco García Iglesias, Gloria Viz Rodríguez, Hugo Alberto Tovar Velasco, Irene Álvarez Rodríguez, Irene García Zuazola, Irene Rivero Calle, Iria Afonso Carrasco, Isabel Ferreirós Vidal, Isabel Lista García, Isabel Rego Lijo, Iván Prieto Gómez, Iván Quintana Cepedal, Jacobo Pardo Seco, Jesús Eirís Puñal, José Gómez Rial, José Manuel Fernández García, José María Martinón Martínez, Julia Cela Mosquera, Julia García Currás, Julián Montoto Louzao, Lara Martínez Martínez, Laura Navarro Marrón, Lidia Piñeiro Rodríguez, Lorenzo Redondo Collazo, Lúa Castelo Martínez, Lucía Company Arciniegas, Luis Crego Rodríguez, Luisa García Vicente, Manuel Vázquez Donsión, María Dolores Martínez García, María Elena Gamborino Caramés, María Elena Sobrino Fernández, María José Currás Tuala, María Martínez Leis, María Soledad Vilas Iglesias, María Sol Rodriguez Calvo, María Teresa Autran García, Marina Casas Pérez, Marta Aldonza Torres, Marta Bouzón Alejandro, Marta Lendoiro Fuentes, Miriam Ben García, Miriam Cebey López, Montserrat López Franco, Nour El Zahraa Mallah, Narmeen Mallah, Natalia García Sánchez, Natalia Vieito Perez, Patricia Regueiro Casuso, Ricardo Suárez Camacho, Rita García Fernández, Rita Varela Estévez, Rosaura Picáns Leis, Ruth Barral Arca, Sandra Carnota Antonio, Sandra Viz Lasheras, Sara Pischedda, Sara Rey Vázquez, Sonia Marcos Alonso, Sonia Serén Fernández, Susana Rey García, Vanesa Álvarez Iglesias, Victoria Redondo Cervantes, Vanesa Álvarez Iglesias, Wiktor Dominik Nowak, Xabier Bello Paderne, Xabier Mazaira López

### Nursing volunteers (alphabetical order)

Alejandra Fernández Méndez, Ana Isabel Abadín Campaña, Ana María León Caamaño, Ana María Buide Illobre, Ángeles Mera Cores, Carmen Nieves Vastro, Carolina Suarez Crego, Concepción Rey Iglesias, Cristina Candal Regueira, Dolores Barreiro Puente, Elvira Rodríguez Rodríguez, Eugenia González Budiño, Eva Rey Álvarez, Fernando Rodríguez Gerpe, Gemma Albela Silva, Isabel Castro Pérez, Isabel Domínguez Ríos, José Ángel Fernández de la Iglesia, José Cruces Vázquez, José Luis Cambeiro Quintela, José Ramón Magariños Iglesias, Julia Rey Brandariz, Julio Abel Fernández López, Luisa García Vicente, Manuel González Lito, Manuel González Lijó, Manuela Pérez Rivas, Margarita Turnes Paredes, María Aurora Méndez López, María Begoña Tomé Arufe, María Campos Torres, María del Carmen Baloira Nogueira, María del Carmen García juan, María Esther Moricosa García, María Luz Chao Jarel, María Martínez Leis, María Mercedes Jiménez Santos, María Salomé Buide Illobre, María Victoria López Pereira, Mercedes Jorge González, Mercedes Isolina Rodríguez Rodríguez, Miren Payo Puente, Natalia Carter Domínguez, Olga María Reyes González, Pilar Mera Rodríguez, Purificación Sebio Brandariz, Salomé Quintáns lago, Yolanda Rodríguez Taboada, María Pereira Grau.

### Other volunteers (alphabetical order)

Alba Arias Gómez, Alejandro Moreno Díaz, Ana Arca Marán, Astro González Guirado, Brais García Iglesias, Carlos Sánchez Rubín, Carmen Otero de Andrés, Clara Pérez Errazquin Barrera, Claudia Rey Posse, Cristina Rojas García, Eduardo Xavier Giménez Bargiela, Elena Gloria Morales García, Fabio Izquierdo García Escribano, Gabriel Guisande García, Jaime López Martín, Lara Pais Ramiro, Lucía Rico Montero, Luís Estévez Martínez, Manuel Estévez Casal, María Aránzazu Palomino Caño, María Rubio Valdés, Marisol Nogales Benítez, Miryam Tilve Pérez, Nuria Villar Muiños, Pablo Del Cerro Rodríguez, Pablo Pozuelo Martínez Cardeñoso, Salma Ouahabi El Ouahabi, Santiago Vázquez Calvache

## Notes

### Competing Interest Statement

The authors have declared no competing interest.

## References

1. Association AP: Diagnostic and statistical manual of mental disorders (5th ed., text rev.; DSM-5-TR): American Psychiatric Publishing; 2022.

2. Kamp-Becker I: Autism spectrum disorder in ICD-11-a critical reflection of its possible impact on clinical practice and research. Mol Psychiatry 2024, 29(3):633–638.

3. Organization WH: International Classification of Diseases, Eleventh Revision (ICD-11). 2021.

4. Zeidan J, Fombonne E, Scorah J, Ibrahim A, Durkin MS, Saxena S, Yusuf A, Shih A, Elsabbagh M: Global prevalence of autism: A systematic review update. Autism Res 2022, 15(5):778–790.

5. The Prevalence of Autism Spectrum Disorder in Europe [www.intechopen.com]

6. Volkmar F, Siegel M, Woodbury-Smith M, King B, McCracken J, State M, American Academy of C, Adolescent Psychiatry Committee on Quality I: Practice parameter for the assessment and treatment of children and adolescents with autism spectrum disorder. J Am Acad Child Adolesc Psychiatry 2014, 53(2):237–257.

7. Delobel-Ayoub M, Saemundsen E, Gissler M, Ego A, Moilanen I, Ebeling H, Rafnsson V, Klapouszczak D, Thorsteinsson E, Arnaldsdottir KM et al: Prevalence of Autism Spectrum Disorder in 7-9-Year-Old Children in Denmark, Finland, France and Iceland: A Population-Based Registries Approach Within the ASDEU Project. J Autism Dev Disord 2020, 50(3):949–959.

8. Volkmar FR, Cohen DJ, Bregman JD, Hooks MY, Stevenson JM: An examination of social typologies in autism. J Am Acad Child Adolesc Psychiatry 1989, 28(1):82–86.

9. Ukkola-Vuoti L, Kanduri C, Oikkonen J, Buck G, Blancher C, Raijas P, Karma K, Lahdesmaki H, Jarvela I: Genome-wide copy number variation analysis in extended families and unrelated individuals characterized for musical aptitude and creativity in music. PLoS One 2013, 8(2):e56356.

10. Muth A, Honekopp J, Falter CM: Visuo-spatial performance in autism: a meta-analysis. J Autism Dev Disord 2014, 44(12):3245–3263.

11. Remington A, Fairnie J: A sound advantage: Increased auditory capacity in autism. Cognition 2017, 166:459–465.

12. Samson F, Mottron L, Soulieres I, Zeffiro TA: Enhanced visual functioning in autism: an ALE meta-analysis. Hum Brain Mapp 2012, 33(7):1553–1581.

13. Mazurek MO, Lu F, Symecko H, Butter E, Bing NM, Hundley RJ, Poulsen M, Kanne SM, Macklin EA, Handen BL: A Prospective Study of the Concordance of DSM-IV and DSM-5 Diagnostic Criteria for Autism Spectrum Disorder. J Autism Dev Disord 2017, 47(9):2783–2794.

14. Kielinen M, Rantala H, Timonen E, Linna SL, Moilanen I: Associated medical disorders and disabilities in children with autistic disorder: a population-based study. Autism 2004, 8(1):49–60.

15. Soke GN, Maenner MJ, Christensen D, Kurzius-Spencer M, Schieve LA: Prevalence of Co-occurring Medical and Behavioral Conditions/Symptoms Among 4- and 8-Year-Old Children with Autism Spectrum Disorder in Selected Areas of the United States in 2010. J Autism Dev Disord 2018, 48(8):2663–2676.

16. Seltzer MM, Shattuck P, Abbeduto L, Greenberg JS: Trajectory of development in adolescents and adults with autism. Ment Retard Dev Disabil Res Rev 2004, 10(4):234–247.

17. Stoodley CJ, D’Mello AM, Ellegood J, Jakkamsetti V, Liu P, Nebel MB, Gibson JM, Kelly E, Meng F, Cano CA et al: Altered cerebellar connectivity in autism and cerebellar-mediated rescue of autism-related behaviors in mice. Nat Neurosci 2017, 20(12):1744–1751.

18. Hazlett HC, Gu H, Munsell BC, Kim SH, Styner M, Wolff JJ, Elison JT, Swanson MR, Zhu H, Botteron KN et al: Early brain development in infants at high risk for autism spectrum disorder. Nature 2017, 542(7641):348–351.

19. McDougle CJ, Erickson CA, Stigler KA, Posey DJ: Neurochemistry in the pathophysiology of autism. J Clin Psychiatry 2005, 66 Suppl 10:9–18.

20. Voineagu I, Wang X, Johnston P, Lowe JK, Tian Y, Horvath S, Mill J, Cantor RM, Blencowe BJ, Geschwind DH: Transcriptomic analysis of autistic brain reveals convergent molecular pathology. Nature 2011, 474(7351):380–384.

21. Zoghbi HY: Postnatal neurodevelopmental disorders: meeting at the synapse? Science 2003, 302(5646):826-830.

22. Bjørk M, Riedel B, Spigset O, Veiby G, Kolstad E, Daltveit AK, Gilhus NE: Association of Folic Acid Supplementation During Pregnancy With the Risk of Autistic Traits in Children Exposed to Antiepileptic Drugs In Utero. JAMA Neurol 2018, 75(2):160–168.

23. Jash S, Sharma S: In utero immune programming of autism spectrum disorder (ASD). Hum Immunol 2021, 82(5):379–384.

24. Zatorre R, McGill J: Music, the food of neuroscience? Nature 2005, 434(7031):312-315.

25. Kanduri C, Raijas P, Ahvenainen M, Philips AK, Ukkola-Vuoti L, Lahdesmaki H, Jarvela I: The effect of listening to music on human transcriptome. PeerJ 2015, 3:e830.

26. Association AMT: Music Therapy Fact Sheets & Bibliographies. In.; 2014.

27. Goris J, Brass M, Cambier C, Delplanque J, Wiersema JR, Braem S: The Relation Between Preference for Predictability and Autistic Traits. Autism Res 2020, 13(7):1144–1154.

28. Cook A, Ogden J, Winstone N: The impact of a school-based musical contact intervention on prosocial attitudes, emotions and behaviours: A pilot trial with autistic and neurotypical children. Autism 2019, 23(4):933–942.

29. Lense MD, Beck S, Liu C, Pfeiffer R, Diaz N, Lynch M, Goodman N, Summers A, Fisher MH: Parents, Peers, and Musical Play: Integrated Parent-Child Music Class Program Supports Community Participation and Well-Being for Families of Children With and Without Autism Spectrum Disorder. Front Psychol 2020, 11:555717.

30. Domes G, Heinrichs M, Michel A, Berger C, Herpertz SC: Oxytocin improves “mind-reading” in humans. Biol Psychiatry 2007, 61(6):731–733.

31. Israel S, Lerer E, Shalev I, Uzefovsky F, Reibold M, Bachner-Melman R, Granot R, Bornstein G, Knafo A, Yirmiya N et al: Molecular genetic studies of the arginine vasopressin 1a receptor (AVPR1a) and the oxytocin receptor (OXTR) in human behaviour: from autism to altruism with some notes in between. Prog Brain Res 2008, 170:435–449.

32. Sharda M, Tuerk C, Chowdhury R, Jamey K, Foster N, Custo-Blanch M, Tan M, Nadig A, Hyde K: Music improves social communication and auditory-motor connectivity in children with autism. Transl Psychiatry 2018, 8(1):231.

33. Sukhodolsky DG, Scahill L, Gadow KD, Arnold LE, Aman MG, McDougle CJ, McCracken JT, Tierney E, Williams White S, Lecavalier L et al: Parent-rated anxiety symptoms in children with pervasive developmental disorders: frequency and association with core autism symptoms and cognitive functioning. J Abnorm Child Psychol 2008, 36(1):117–128.

34. Wolff JJ, Symons FJ: An evaluation of multi-component exposure treatment of needle phobia in an adult with autism and intellectual disability. J Appl Res Intellect Disabil 2013, 26(4):344–348.

35. Meindl JN, Saba S, Gray M, Stuebing L, Jarvis A: Reducing blood draw phobia in an adult with autism spectrum disorder using low-cost virtual reality exposure therapy. J Appl Res Intellect Disabil 2019, 32(6):1446–1452.

36. Farah R, Haraty H, Salame Z, Fares Y, Ojcius DM, Said Sadier N: Salivary biomarkers for the diagnosis and monitoring of neurological diseases. Biomed J 2018, 41(2):63–87.

37. Engeland CG, Bosch JA, Rohleder N: Salivary Biomarkers in Psychoneuroimmunology. Curr Opin Behav Sci 2019, 28:58–65.

38. Nakamura T, Matsui M, Uchida K, Futatsugi A, Kusakawa S, Matsumoto N, Nakamura K, Manabe T, Taketo MM, Mikoshiba K: M(3) muscarinic acetylcholine receptor plays a critical role in parasympathetic control of salivation in mice. J Physiol 2004, 558(Pt 2):561–575.

39. Murai S, Saito H, Masuda Y, Itoh T, Kawaguchi T: Sex-dependent differences in the concentrations of the principal neurotransmitters, noradrenaline and acetylcholine, in the three major salivary glands of mice. Arch Oral Biol 1998, 43(1):9–14.

40. Proctor GB, Carpenter GH: Regulation of salivary gland function by autonomic nerves. Auton Neurosci 2007, 133(1):3–18.

41. Sansores-Espana LD, Melgar-Rodriguez S, Olivares-Sagredo K, Cafferata EA, Martinez-Aguilar VM, Vernal R, Paula-Lima AC, Diaz-Zuniga J: Oral-Gut-Brain Axis in Experimental Models of Periodontitis: Associating Gut Dysbiosis With Neurodegenerative Diseases. Front Aging 2021, 2:781582.

42. Hicks SD, Rajan AT, Wagner KE, Barns S, Carpenter RL, Middleton FA: Validation of a Salivary RNA Test for Childhood Autism Spectrum Disorder. Frontiers in genetics 2018, 9:534.

43. Ragusa M, Santagati M, Mirabella F, Lauretta G, Cirnigliaro M, Brex D, Barbagallo C, Domini CN, Gulisano M, Barone R et al: Potential Associations Among Alteration of Salivary miRNAs, Saliva Microbiome Structure, and Cognitive Impairments in Autistic Children. International journal of molecular sciences 2020, 21(17).

44. Benach JL, Li E, McGovern MM: A microbial association with autism. mBio 2012, 3(1).

45. Bakker-Huvenaars MJ, Greven CU, Herpers P, Wiegers E, Jansen A, van der Steen R, van Herwaarden AE, Baanders AN, Nijhof KS, Scheepers F et al: Saliva oxytocin, cortisol, and testosterone levels in adolescent boys with autism spectrum disorder, oppositional defiant disorder/conduct disorder and typically developing individuals. Eur Neuropsychopharmacol 2020, 30:87–101.

46. Majewska MD, Hill M, Urbanowicz E, Rok-Bujko P, Bienkowski P, Namyslowska I, Mierzejewski P: Marked elevation of adrenal steroids, especially androgens, in saliva of prepubertal autistic children. Eur Child Adolesc Psychiatry 2014, 23(6):485–498.

47. John S, Jaeggi AV: Oxytocin levels tend to be lower in autistic children: A meta-analysis of 31 studies. Autism 2021, 25(8):2152–2161.

48. Yatawara CJ, Einfeld SL, Hickie IB, Davenport TA, Guastella AJ: The effect of oxytocin nasal spray on social interaction deficits observed in young children with autism: a randomized clinical crossover trial. Mol Psychiatry 2016, 21(9):1225–1231.

49. Parker KJ, Oztan O, Libove RA, Sumiyoshi RD, Jackson LP, Karhson DS, Summers JE, Hinman KE, Motonaga KS, Phillips JM et al: Intranasal oxytocin treatment for social deficits and biomarkers of response in children with autism. Proc Natl Acad Sci U S A 2017, 114(30):8119–8124.

50. Munesue T, Nakamura H, Kikuchi M, Miura Y, Takeuchi N, Anme T, Nanba E, Adachi K, Tsubouchi K, Sai Y et al: Oxytocin for Male Subjects with Autism Spectrum Disorder and Comorbid Intellectual Disabilities: A Randomized Pilot Study. Front Psychiatry 2016, 7:2.

51. Anagnostou E, Soorya L, Chaplin W, Bartz J, Halpern D, Wasserman S, Wang AT, Pepa L, Tanel N, Kushki A et al: Intranasal oxytocin versus placebo in the treatment of adults with autism spectrum disorders: a randomized controlled trial. Mol Autism 2012, 3(1):16.

52. Rylander-Rudqvist T, Hakansson N, Tybring G, Wolk A: Quality and quantity of saliva DNA obtained from the self-administrated oragene method--a pilot study on the cohort of Swedish men. Cancer epidemiology, biomarkers & prevention : a publication of the American Association for Cancer Research, cosponsored by the American Society of Preventive Oncology 2006, 15(9):1742–1745.

53. Spielmann N, Ilsley D, Gu J, Lea K, Brockman J, Heater S, Setterquist R, Wong DT: The human salivary RNA transcriptome revealed by massively parallel sequencing. Clin Chem 2012, 58(9):1314–1321.

54. Gosch A, Banemann R, Dorum G, Haas C, Hadrys T, Haenggi N, Kulstein G, Neubauer J, Courts C: Spitting in the wind?-The challenges of RNA sequencing for biomarker discovery from saliva. Int J Legal Med 2024, 138(2):401–412.

55. Christodoulides N, Mohanty S, Miller CS, Langub MC, Floriano PN, Dharshan P, Ali MF, Bernard B, Romanovicz D, Anslyn E et al: Application of microchip assay system for the measurement of C-reactive protein in human saliva. Lab Chip 2005, 5(3):261–269.

56. Gröschl M, Wagner R, Rauh M, Dorr HG: Stability of salivary steroids: the influences of storage, food and dental care. Steroids 2001, 66(10):737–741.

57. Pfaffe T, Cooper-White J, Beyerlein P, Kostner K, Punyadeera C: Diagnostic potential of saliva: current state and future applications. Clin Chem 2011, 57(5):675–687.

58. Gomez-Carballa A, Navarro L, Pardo-Seco J, Bello X, Pischedda S, Viz-Lasheras S, Camino-Mera A, Curras MJ, Ferreiros I, Mallah N et al: Music compensates for altered gene expression in age-related cognitive disorders. Scientific reports 2023, 13(1):21259.

59. Navarro L, Gomez-Carballa A, Pischedda S, Montoto-Louzao J, Viz-Lasheras S, Camino-Mera A, Hinault T, Martinon-Torres F, Salas A: Sensogenomics of music and Alzheimer’s disease: An interdisciplinary view from neuroscience, transcriptomics, and epigenomics. Front Aging Neurosci 2023, 15:1063536.

60. Navarro L, Martinon-Torres F, Salas A: Sensogenomics and the Biological Background Underlying Musical Stimuli: Perspectives for a New Era of Musical Research. Genes (Basel) 2021, 12(9).

61. Bolger AM, Lohse M, Usadel B: Trimmomatic: a flexible trimmer for Illumina sequence data. Bioinformatics 2014, 30(15):2114–2120.

62. Dobin A, Davis CA, Schlesinger F, Drenkow J, Zaleski C, Jha S, Batut P, Chaisson M, Gingeras TR: STAR: ultrafast universal RNA-seq aligner. Bioinformatics 2013, 29(1):15–21.

63. Putri GH, Anders S, Pyl PT, Pimanda JE, Zanini F: Analysing high-throughput sequencing data in Python with HTSeq 2.0. Bioinformatics 2022, 38(10):2943–2945.

64. Love MI, Huber W, Anders S: Moderated estimation of fold change and dispersion for RNA-seq data with DESeq2. Genome Biol 2014, 15(12):550.

65. Wickham H: ggplot2: Elegant Graphics for Data Analysis: Springer-Verlag New York; 2016.

66. Gu Z: Complex heatmap visualization. Imeta 2022, 1(3):e43.

67. Blighe K, Rana S, Lewis M: EnhancedVolcano: Publication-ready volcano plots with enhanced colouring and labeling. version 180, R package. In.; 2020.

68. Hänzelmann S, Castelo R, Guinney J: GSVA: gene set variation analysis for microarray and RNA-seq data. BMC Bioinformatics 2013, 14:7.

69. Ritchie ME, Phipson B, Wu D, Hu Y, Law CW, Shi W, Smyth GK: *limma* powers differential expression analyses for RNA-sequencing and microarray studies. Nucleic Acids Res 2015, 43(7):e47.

70. Wu T, Hu E, Xu S, Chen M, Guo P, Dai Z, Feng T, Zhou L, Tang W, Zhan L et al: clusterProfiler 4.0: A universal enrichment tool for interpreting omics data. Innovation (N Y) 2021, 2(3):100141.

71. Yu G: Thirteen years of clusterProfiler. Innovation (Camb) 2024, 5(6):100722.

72. Carlson M: org.Hs.eg.db: Genome wide annotation for Human. In.; 2023.

73. Pagès H, Carlson M, Falcon S, Li N: AnnotationDbi: Manipulation of SQLite-based annotations in Bioconductor. In.; 2024.

74. Abrahams BS, Arking DE, Campbell DB, Mefford HC, Morrow EM, Weiss LA, Menashe I, Wadkins T, Banerjee-Basu S, Packer A: SFARI Gene 2.0: a community-driven knowledgebase for the autism spectrum disorders (ASDs). Mol Autism 2013, 4(1):36.

75. Langfelder P, Horvath S: WGCNA: an R package for weighted correlation network analysis. BMC Bioinformatics 2008, 9:559.

76. Ounit R, Lonardi S: Higher classification sensitivity of short metagenomic reads with CLARK-S. Bioinformatics 2016, 32(24):3823–3825.

77. Ounit R, Wanamaker S, Close TJ, Lonardi S: CLARK: fast and accurate classification of metagenomic and genomic sequences using discriminative k-mers. BMC Genomics 2015, 16(1):236.

78. waffle: Create Waffle Chart Visualizations. In CRAN: Contributed Packages [10.32614/CRAN.package.waffle]

79. Sanchez-Tena S, Cubillos-Rojas M, Schneider T, Rosa JL: Functional and pathological relevance of HERC family proteins: a decade later. Cellular and molecular life sciences : CMLS 2016, 73(10):1955–1968.

80. Oudshoorn D, van Boheemen S, Sanchez-Aparicio MT, Rajsbaum R, Garcia-Sastre A, Versteeg GA: HERC6 is the main E3 ligase for global ISG15 conjugation in mouse cells. PLoS One 2012, 7(1):e29870.

81. Moretto E, Longatti A, Murru L, Chamma I, Sessa A, Zapata J, Hosy E, Sainlos M, Saint-Pol J, Rubinstein E et al: TSPAN5 Enriched Microdomains Provide a Platform for Dendritic Spine Maturation through Neuroligin-1 Clustering. Cell Rep 2019, 29(5):1130–1146 e1138.

82. Rawsthorne H, Calahorro F, Feist E, Holden-Dye L, O’Connor V, Dillon J: Neuroligin dependence of social behaviour in Caenorhabditis elegans provides a model to investigate an autism-associated gene. Hum Mol Genet 2021, 29(21):3546–3553.

83. Ghiretti AE, Paradis S: The GTPase Rem2 regulates synapse development and dendritic morphology. Dev Neurobiol 2011, 71(5):374–389.

84. Rubenstein JL, Merzenich MM: Model of autism: increased ratio of excitation/inhibition in key neural systems. Genes Brain Behav 2003, 2(5):255–267.

85. Belichenko PV, Kleschevnikov AM, Masliah E, Wu C, Takimoto-Kimura R, Salehi A, Mobley WC: Excitatory-inhibitory relationship in the fascia dentata in the Ts65Dn mouse model of Down syndrome. The Journal of comparative neurology 2009, 512(4):453–466.

86. Li X, Chauhan A, Sheikh AM, Patil S, Chauhan V, Li XM, Ji L, Brown T, Malik M: Elevated immune response in the brain of autistic patients. J Neuroimmunol 2009, 207(1-2):111–116.

87. Robinson-Agramonte MLA, Noris Garcia E, Fraga Guerra J, Vega Hurtado Y, Antonucci N, Semprun-Hernandez N, Schultz S, Siniscalco D: Immune Dysregulation in Autism Spectrum Disorder: What Do We Know about It? International journal of molecular sciences 2022, 23(6).

88. Wei H, Zou H, Sheikh AM, Malik M, Dobkin C, Brown WT, Li X: IL-6 is increased in the cerebellum of autistic brain and alters neural cell adhesion, migration and synaptic formation. J Neuroinflammation 2011, 8:52.

89. Shin SM, Zhang N, Hansen J, Gerges NZ, Pak DT, Sheng M, Lee SH: GKAP orchestrates activity-dependent postsynaptic protein remodeling and homeostatic scaling. Nat Neurosci 2012, 15(12):1655–1666.

90. Chen Q, Zhu YC, Yu J, Miao S, Zheng J, Xu L, Zhou Y, Li D, Zhang C, Tao J et al: CDKL5, a protein associated with rett syndrome, regulates neuronal morphogenesis via Rac1 signaling. J Neurosci 2010, 30(38):12777–12786.

91. Maracci C, Motta S, Romagnoli A, Costantino M, Perego P, Di Marino D: The mTOR/4E-BP1/eIF4E Signalling Pathway as a Source of Cancer Drug Targets. Curr Med Chem 2022, 29(20):3501–3529.

92. Sun CY, van Koningsbruggen S, Long SW, Straasheijm K, Klooster R, Jones TI, Bellini M, Levesque L, Brieher WM, van der Maarel SM et al: Facioscapulohumeral muscular dystrophy region gene 1 is a dynamic RNA-associated and actin-bundling protein. Journal of molecular biology 2011, 411(2):397–416.

93. Kim N, Kim KH, Lim WJ, Kim J, Kim SA, Yoo HJ: Whole Exome Sequencing Identifies Novel De Novo Variants Interacting with Six Gene Networks in Autism Spectrum Disorder. Genes (Basel) 2020, 12(1).

94. Guo D, Yang X, Shi L: Rho GTPase Regulators and Effectors in Autism Spectrum Disorders: Animal Models and Insights for Therapeutics. Cells 2020, 9(4).

95. Arrazola Sastre A, Luque Montoro M, Galvez-Martin P, Lacerda HM, Lucia AM, Llavero F, Zugaza JL: Small GTPases of the Ras and Rho Families Switch on/off Signaling Pathways in Neurodegenerative Diseases. International journal of molecular sciences 2020, 21(17).

96. Zeidan-Chulia F, de Oliveira BN, Casanova MF, Casanova EL, Noda M, Salmina AB, Verkhratsky A: Up-Regulation of Oligodendrocyte Lineage Markers in the Cerebellum of Autistic Patients: Evidence from Network Analysis of Gene Expression. Mol Neurobiol 2016, 53(6):4019–4025.

97. Ricci S, Businaro R, Ippoliti F, Lo Vasco VR, Massoni F, Onofri E, Troili GM, Pontecorvi V, Morelli M, Rapp Ricciardi M et al: Altered cytokine and BDNF levels in autism spectrum disorder. Neurotox Res 2013, 24(4):491–501.

98. Careaga M, Rogers S, Hansen RL, Amaral DG, Van de Water J, Ashwood P: Immune Endophenotypes in Children With Autism Spectrum Disorder. Biol Psychiatry 2017, 81(5):434–441.

99. Saghazadeh A, Ataeinia B, Keynejad K, Abdolalizadeh A, Hirbod-Mobarakeh A, Rezaei N: A meta-analysis of pro-inflammatory cytokines in autism spectrum disorders: Effects of age, gender, and latitude. J Psychiatr Res 2019, 115:90–102.

100. Fujita E, Dai H, Tanabe Y, Zhiling Y, Yamagata T, Miyakawa T, Tanokura M, Momoi MY, Momoi T: Autism spectrum disorder is related to endoplasmic reticulum stress induced by mutations in the synaptic cell adhesion molecule, CADM1. Cell Death Dis 2010, 1(6):e47.

101. Kawada K, Mimori S: Implication of Endoplasmic Reticulum Stress in Autism Spectrum Disorder. Neurochem Res 2018, 43(1):147–152.

102. Crider A, Ahmed AO, Pillai A: Altered Expression of Endoplasmic Reticulum Stress-Related Genes in the Middle Frontal Cortex of Subjects with Autism Spectrum Disorder. Mol Neuropsychiatry 2017, 3(2):85–91.

103. Wong GC, Montgomery JM, Taylor MW: The Gut-Microbiota-Brain Axis in Autism Spectrum Disorder, vol. 8. Brisbane (AU): Exon Publications; 2021.

104. D’Angelo E, Fiori F, Ferraro GA, Tessitore A, Nazzaro L, Serpico R, Contaldo M: Autism Spectrum Disorder, Oral Implications, and Oral Microbiota. Children (Basel) 2025, 12(3).

105. Coronas R, Zara G, Gallo A, Rocchetti G, Lapris M, Petretto GL, Zara S, Fancello F, Mannazzu I: Propionibacteria as promising tools for the production of pro-bioactive scotta: a proof-of-concept study. Front Microbiol 2023, 14:1223741.

106. Sato-Suzuki Y, Washio J, Wicaksono DP, Sato T, Fukumoto S, Takahashi N: Nitrite-producing oral microbiome in adults and children. Scientific reports 2020, 10(1):16652.

107. Srikantha P, Mohajeri MH: The Possible Role of the Microbiota-Gut-Brain-Axis in Autism Spectrum Disorder. International journal of molecular sciences 2019, 20(9).

108. Li Q, Zhou JM: The microbiota-gut-brain axis and its potential therapeutic role in autism spectrum disorder. Neuroscience 2016, 324:131–139.

109. El-Ansary A, Bhat RS, Al-Daihan S, Al Dbass AM: The neurotoxic effects of ampicillin-associated gut bacterial imbalances compared to those of orally administered propionic acid in the etiology of persistent autistic features in rat pups: effects of various dietary regimens. Gut Pathog 2015, 7:7.

110. El-Ansary AK, Ben Bacha A, Kotb M: Etiology of autistic features: the persisting neurotoxic effects of propionic acid. J Neuroinflammation 2012, 9:74.

111. Feng P, Zhao S, Zhang Y, Li E: A review of probiotics in the treatment of autism spectrum disorders: Perspectives from the gut-brain axis. Front Microbiol 2023, 14:1123462.

112. Frye RE, Slattery J, MacFabe DF, Allen-Vercoe E, Parker W, Rodakis J, Adams JB, Krajmalnik-Brown R, Bolte E, Kahler S et al: Approaches to studying and manipulating the enteric microbiome to improve autism symptoms. Microb Ecol Health Dis 2015, 26:26878.

