## Supplementary figures and images for "Decoding the peripheral transcriptomic and meta-genomic response to music in Autism Spectrum Disorder *via* saliva-based RNA sequencing"

### Figure S1

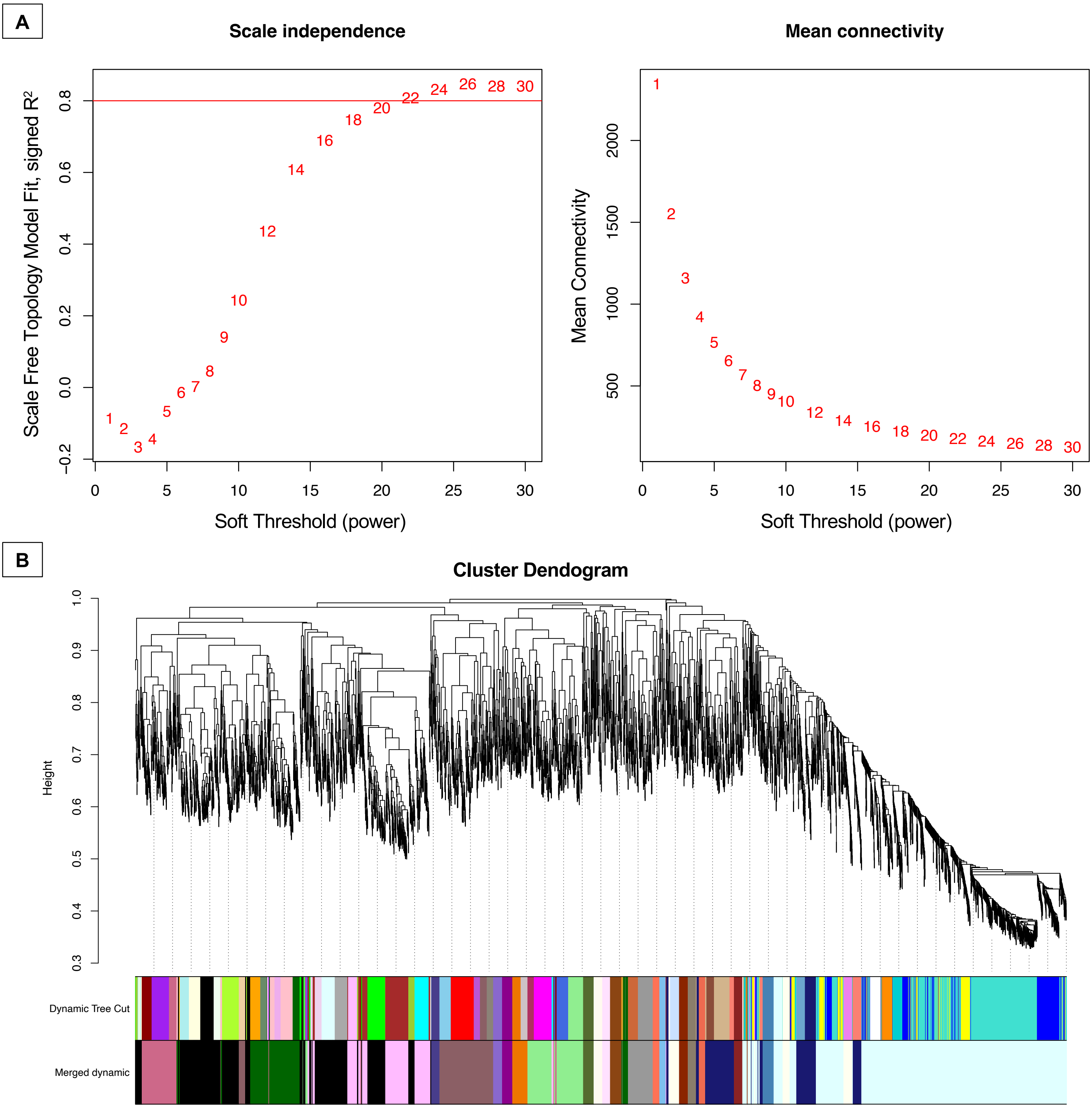

### Figure S2

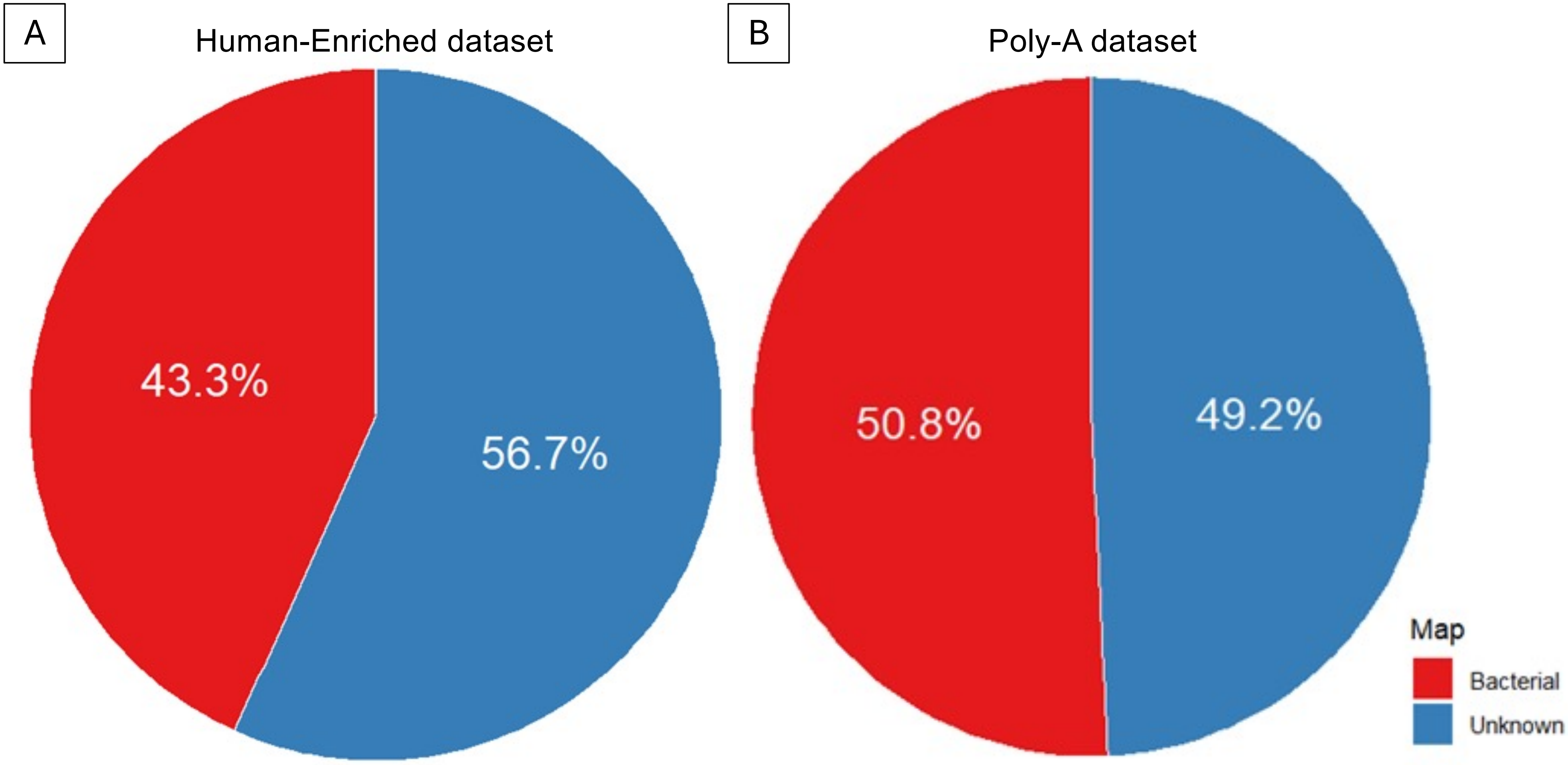
